# Leveraging clinical sleep data across multiple pediatric cohorts for insights into neurodevelopment: the Retrospective Analysis of Sleep in Pediatric (RASP) cohorts study

**DOI:** 10.1101/2024.10.04.616563

**Authors:** Naihua N. Gong, Aditya Mahat, Samya Ahmad, Daniel Glaze, Mirjana Maletic-Savatic, Matthew McGinley, Anne Marie Morse, Alcibiades J. Rodriguez, Audrey Thurm, Susan Redline, Kiran Maski, Shaun Purcell, Ashura Buckley

## Abstract

Sleep disturbances are prominent across neurodevelopmental disorders (NDDs) and may reflect specific abnormalities in brain development and function. Overnight polysomnography (PSG) allows for detailed investigation of sleep architecture, offering a unique window into neurocircuit function. A better understanding of sleep in NDDs compared to typically developing children could therefore define mechanisms underlying abnormal development in NDDs and provide avenues for the development of therapeutic interventions to improve sleep quality and developmental outcomes. Here, we introduce and characterize a collection of 1527 pediatric overnight PSGs across five different sites. We first developed an automated stager trained on independent pediatric sleep data, which yielded better performance compared to a stager trained on adults. Using consistent staging across cohorts, we derived a panel of EEG micro-architectural features. This unbiased approach replicated broad trajectories previously described in typically developing sleep architecture. Further, we found sleep architecture disruptions in children with Down’s Syndrome (DS) that were consistent across independent cohorts. Finally, we built and evaluated a model to predict age from sleep EEG metrics, which recapitulated our previous findings of younger predicted brain age in children with DS. Altogether, by creating a resource pooled from existing clinical data we expanded the available datasets and computational resources to study sleep in pediatric populations, specifically towards a better understanding of sleep in NDDs. This Retrospective Analysis of Sleep in Pediatric (RASP) cohorts dataset, including staging annotation derived from our automated stager, will be deposited at https://sleepdata.org.

**Statement of significance:** We introduce the RASP cohorts, a collection of 1527 clinical pediatric overnight polysomnographies that includes typically developing and neurodevelopmental disorder cases. As a first step towards addressing the analytic bottleneck inherent in manual sleep staging, we developed and validated a pediatric-specific sleep stager. Leveraging the retrospective RASP cohorts dataset, we redemonstrated known developmental trajectories in sleep architecture. To summarize changes in brain function reflected in sleep, we developed a model to predict brain age from sleep measures. We recapitulate younger predicted age in RASP Down’s Syndrome cases. This work not only enhances our understanding of sleep disturbances in NDDs, but also provides a valuable resource for future research and underscores the utility of existing clinical polysomnography studies.

## Introduction

Sleep in early life is characterized by increased duration and distinct architecture compared to sleep in maturity [1–5]. Across species, unique juvenile sleep state dynamics are thought to be essential for neurodevelopment, particularly during critical periods of enhanced plasticity [6–11]. In the last decade, the prevalence of disrupted sleep in the US pediatric population has increased, with less than half of children and adolescents getting the recommended amount of nightly sleep [12–15]. While the effects of sleep on cognitive functions, mood, and emotional regulation in adults are well-documented, there is emerging evidence that sleep disturbances in children and adolescents result in more pronounced negative consequences in these domains [16–18]. Longitudinally, sleep difficulties in early life are associated with later behavioral and emotional dysregulation [19–21]. Inadequate sleep during childhood poses an increased risk to neuropsychological health during sensitive periods of brain maturation and may ultimately affect the trajectory of neurodevelopment. A better understanding of early life sleep and how specific disruptions relate to adverse outcomes is essential for developing effective early interventions.

Neurodevelopmental disorders (NDDs) are disorders of brain development that are highly heterogeneous in presentation but commonly share sleep-wake disturbances as some of the earliest clinical features [22–24]. The higher rate of sleep problems in NDDs suggests a mechanistic link between childhood sleep and brain development. Indeed, shorter sleep duration is associated with increased severity of social impairments and other behavioral symptoms in ASD [25]. However, sleep studies in children with NDDs typically rely on parent reporting of metrics such as time spent in bed and night time awakenings [26], which may inaccurately reflect nuanced disturbances in sleep architecture. Quantifying sleep signatures beyond sleep duration and subjective measures of sleep quality is critical for understanding how sleep disruptions might also be variable across and within NDDs.

Alterations in sleep architecture offer a unique lens into neural activity in NDDs and may be useful for implicating specific neurocircuit dysfunction within these disorders. For example, sleep spindles, short bursts of neural activity in the 11-15 Hz frequency range that occur in non-rapid eye movement (NREM) sleep, are generated by thalamic activity propagating to higher order cortical layers. Sleep spindle abnormalities in disorders like schizophrenia or dementia reflect thalamocortical dysfunction [27,28]. Objective measures of sleep neurophysiology in the context of NDDs will inform our understanding of the mechanisms underlying early life sleep disruptions in the context of abnormal neurodevelopment and inform our diagnostic and therapeutic approach towards NDDs.

Overnight polysomnography (PSG) studies are the gold standard for studying sleep patterns. The clinical standard for analysis of overnight PSGs is based on manual sleep scoring, in which trained technicians score sleep studies in 30-second epochs, delineating five standard stages based on a limited EEG montage (based on the American Academy of Sleep Medicine manual): wake (W), three stages of NREM (N1, N2, and N3), and rapid eye movement (REM) sleep [29], with special considerations for scoring pediatric studies, in particular in very early life (i.e., the first two months) when sleep stages comprise active, quiet, and indeterminate sleep [30]. This is a time-consuming process in large-scale sleep studies, warranting automation of PSG analysis to increase the speed and consistency of data analysis. These approaches have been utilized in general populations with models trained primarily on adult sleep studies with success [31,32]. However, the distinct sleep architecture of early life may diminish performance of generic models in predicting sleep stages in pediatric populations [33].

From staged PSGs, individual differences in sleep EEG micro-architecture can be quantified through lme-frequency analysis, including stage-specific characterization of activity (spectral power) and global patterns of functional interaction (coherence). Furthermore, staged PSGs facilitate the identification of transient events during NREM, including spindles and slow oscillations, characterized in terms of their occurrence, morphology and temporal coupling. In the context of a pediatric sample, one approach to summarizing sleep metrics is a composite that optimally tracks with development as indexed by chronological age. Metrics from overnight PSGs can be used in predictive models of “brain age” [28,34]. Calculated predicted age difference (PAD) (actual age – predicted brain age) can be interpreted as a measure of overall brain function. In adults with neuropsychiatric disease, sleep EEG-predicted brain age was older than that of healthy controls [34,35].

In pediatric contexts, recent work from our group demonstrated the transferability of sleep EEG-derived findings across different pediatric clinical and research cohorts through an age prediction model trained on pediatric sleep EEG data. Using the Nationwide Children’s Hospital Sleep Databank (NCH-SDB) [36], we characterized macro-as well as microarchitectural changes in sleep in children without NDDs, showing developmental progression in sleep duration and stage, along with changes in spindles and slow oscillations during NREM sleep.

Importantly, we developed a model to predict chronological age from sleep EEGs, which accurately inferred age in another independent pediatric cohort without NDDs. Strikingly, 140 children with Down’s syndrome (DS) aged 2.5 to 17.4 years exhibited younger age predictions compared to age-matched children without NDDs [28]. These findings show pediatric sleep EEG metrics are comparable across independent cohorts and can objectively define whether neurodevelopment is impacted in NDDs, making a case for multi-insltulonal collaborative pediatric sleep data collection initiatives.

Here, we expand on pediatric sleep resources with the Retrospective Analysis of Sleep in Pediatric (RASP) cohort, which comprises 1527 overnight PSGs across five study sites, representing 1326 unique individuals. Manual staging was not available for all RASP cohorts.

This, along with the goal to improve standardization of staging across clinical cohorts, motivated us to develop and apply a pediatric automated stager based on extant data in the National Sleep Research Resource (NSRR), a database of PSG studies across the lifespan that contains several pediatric cohorts [36]. Across diagnostic status, we show a stager trained on pediatric data is more accurate at predicting stages in children compared to a model trained primarily on adults. Utilizing our stager, we generated sleep stage annotations for downstream analysis in the RASP cohorts and demonstrate conservation of NREM changes in sleep EEGs across development in the RASP cohorts. We find specific disruptions in sleep architecture in DS but not autism spectrum disorder (ASD) that are conserved in independent cohorts from the NCH-SDB. Finally, expanding on our prior work [28], we built an age prediction model based on a large training dataset of three existing pediatric cohorts. Using data from NCH-SDB, CHAT, and PATS, we identified sleep EEG features that demonstrated robust changes with age. This approach additionally allowed for the identification of cross-cohort artifacts, which we removed from our predictive model. We found a high degree of concordance between reported and predicted age across the RASP cohorts, recapitulating previous findings of younger brain age in DS [28]. This resource expands on existing datasets of pediatric sleep studies and demonstrates broad comparability across PSG study sites that can be leveraged for more precise characterization of sleep in neurodevelopment.

## Results

### Demographics of the RASP cohorts

The collected dataset consisted of 1527 whole night PSGs from 1326 pediatric individuals from five different institutions: Boston Children’s Hospital (BCH), Geisinger Health System (GEIS), National Institute of Mental Health (NIMH), New York University Langone Comprehensive Epilepsy Center-Sleep Center (NYU), and Texas Children’s Hospital (TCH).

Individuals ranged in age from postpartum day 3 to 18 years of age, with 604 female subjects (39.6%), and 509 subjects with a diagnosed NDD (149 or 29.4% of which were female). Given differences in very early life sleep architecture and rapid changes in sleep in early life, we restricted all subsequent analysis to cases 2.5 years and older (N = 1114 recordings from 955 individuals; see **Table 1** for demographics of analyzed cases). For the purposes of this study, we focused on DS and ASD as NDDs of interest. Most patients without NDDs underwent whole night sleep studies for the evaluation of primary sleep-related conditions (see **Supplementary Table 1** for full list of diagnoses).

**Table 1:** RASP recordings demographics and diagnoses, age ≥ 2.5 years.

| Sex | Mean age ± SD | Minimum age | Maximum age | non-NDD | NDD | ASD | DS |
| --- | --- | --- | --- | --- | --- | --- | --- |
| **BCH** |  |  |  |  |  |  |  |
| Female | 3.92 ± 0.8 | 2.50 | 4.92 | 21 | 11 | 0 | 3 |
| Male | 4.08 ± 0.7 | 2.50 | 5.00 | 35 | 30 | 5 | 6 |
| **NIMH\*** |  |  |  |  |  |  |  |
| Female | 5.44 ± 1.8 | 2.76 | 11.36 | 21 | 69 | 47 | 0 |
| Male | 5.33 ± 1.8 | 2.50 | 10.23 | 36 | 218 | 180 | 0 |
| **NYU** |  |  |  |  |  |  |  |
| Female | 7.7 ± 4 | 2.83 | 15.42 | 8 | 6 | 0 | 1 |
| Male | 6.84 ± 4 | 2.67 | 16.83 | 15 | 6 | 2 | 0 |
| **Geisinger** |  |  |  |  |  |  |  |
| Female | 3.5 ± 0.6 | 3.00 | 5.00 | 21 | 3 | 0 | 1 |
| Male | 3.41 ± 0.5 | 3.00 | 4.00 | 18 | 14 | 3 | 4 |
| **TCH** |  |  |  |  |  |  |  |
| Female | 6.46 ± 4.1 | 3.00 | 18.00 | 252 | 10 | 4 | 2 |
| Male | 6.07 ± 3.7 | 3.00 | 19.00 | 296 | 23 | 9 | 3 |
| **Overall** |  |  |  |  |  |  |  |
| Female | 5.91 ± 3.6 | 2.50 | 18.00 | 323 | 101 | 51 | 7 |
| Male | 5.51 ± 3 | 2.50 | 19.00 | 400 | 291 | 199 | 13 |

\* Counts listed are for number of recordings; NIMH cohort included multiple recordings at different time points for certain individuals.

### Pediatric sleep stagers perform beBer on pediatric cases compared to adult stagers

As GEIS and TCH cases did not have available manual sleep staging, we set out to develop and evaluate an automated stager to apply to the entire RASP cohorts. We first asked whether stagers trained on pediatric data perform better than those trained on primarily adult data. We trained two pediatric models (a “reduced” model including only central EEG channels, and a “full” model additionally including frontal and occipital channels; see Methods for more details) and compared staging performance with results from an adult stager. BCH, NYU, and NIMH had available manual staging (we refer to this group as RASP-M from here onwards), which we leveraged to evaluate the performance of our automated stager. In the overall RASP-M cohort, the two pediatric stagers had better performance compared to the adult stager in both typically developing and NDD cases (for full pediatric, reduced pediatric, and adult stagers, overall median kappa coefficients and IQR for 5-class staging were 0.72 and 0.14, 0.72 and 0.15, and 0.67 and 0.17, respectively; for 3-class (NREM/REM/wake) staging, 0.81 and 0.14, 0.80 and 0.15, and 0.75 and 0.21 respectively; **Figure 1A, C**). Within individual RASP-M cohorts, the pediatric stagers performed better compared to the adult stager for both non-NDD and NDD BCH and NIMH cases. In typically developing NYU cases, only the full pediatric stager had better performance compared to the adult stager, and there was no significant difference between stagers in NDD cases (**Figure S1A; Table 2**). We also assessed staging accuracy for epochs at stage transitions, where we expect greater variability in classification and lower average staging performance. Across NDD status, both the full and reduced pediatric stager had greater accuracy for transition-adjacent epochs compared to the adult stager (**Figure S1B; Table 3A**).

**Figure 1:**
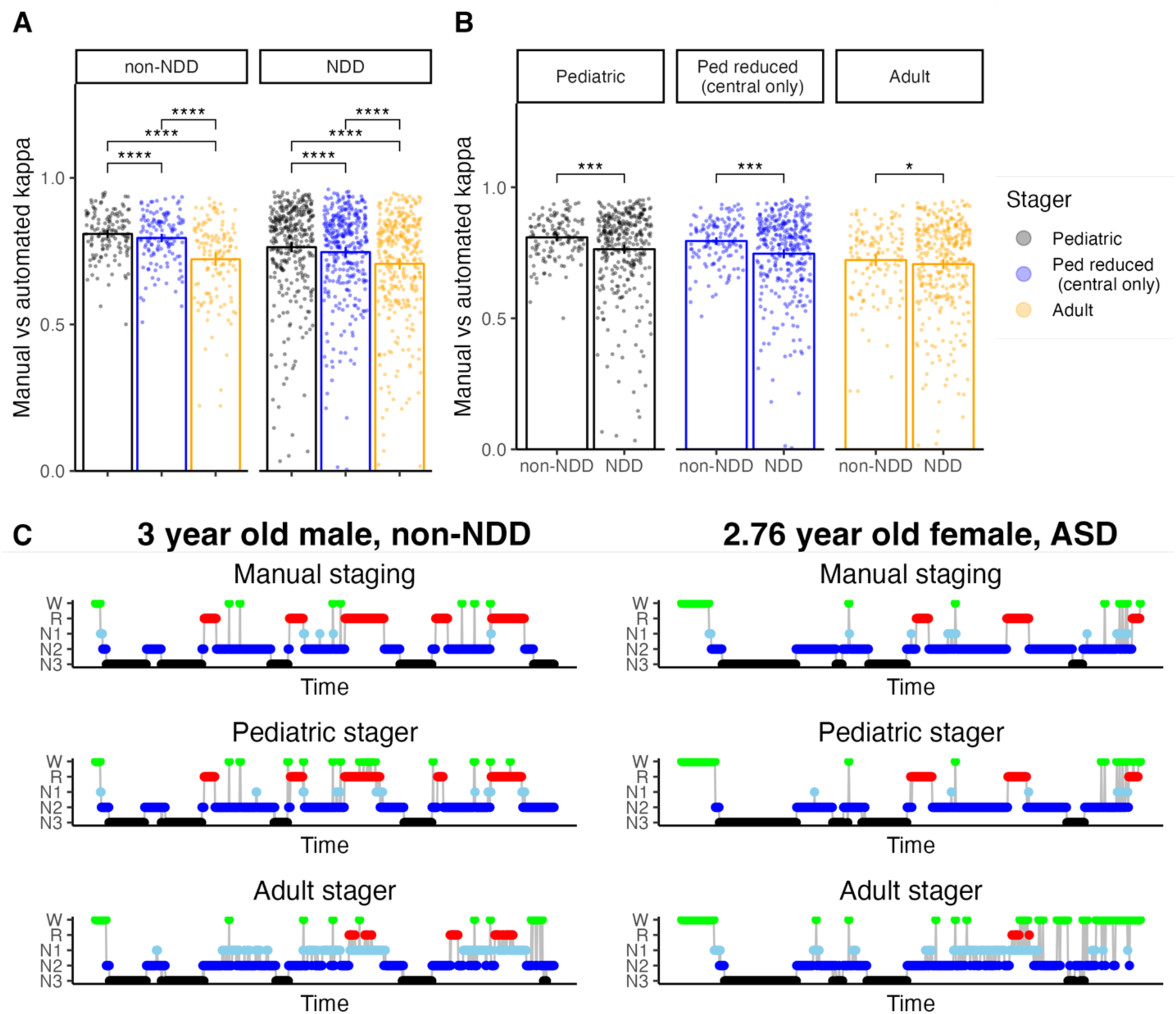
Automated pediatric stagers perform beRer across diagnosSc status in RASP compared to adult stagers. Kappa for 3-class staging between observed vs predicted stage by epoch per individual A) in pediatric stagers (black and blue) and adult stager (yellow), separated by NDD diagnostic status, and B) comparing typical development versus NDD cases across all three automated stagers. C) Example hypnograms comparing manual staging and pediatric vs adult stager in two different individuals from BCH (right) and NIMH (lev). Stalslcal comparisons in A were made using multiple paired T-tests with Bonferroni correction; comparisons in B were made using logistic regression correcting for age, sex, and study site (* p<0.05, ** p < 0.01, *** p < 0.001 for this figure and all subsequent figures).

**Table 2:** Three-stage pediatric vs adult stager performance.

|  | Model | Median kappa | IQR of kappa | Median accuracy | IQR of accuracy | N |
| --- | --- | --- | --- | --- | --- | --- |
| **BCH** |  |  |  |  |  |  |
|  | non-NDD Pediatric | 0.82 | 0.07 | 0.91 | 0.04 | 56 |
|  | non-NDD Ped reduced (central only) | 0.82 | 0.09 | 0.91 | 0.04 | 56 |
|  | non-NDD Adult | 0.73 | 0.20 | 0.88 | 0.08 | 56 |
| **NIMH** |  |  |  |  |  |  |
|  | non-NDD Pediatric | 0.81 | 0.11 | 0.91 | 0.05 | 56 |
|  | non-NDD Ped reduced (central only) | 0.79 | 0.10 | 0.90 | 0.05 | 56 |
|  | non-NDD Adult | 0.75 | 0.13 | 0.89 | 0.06 | 56 |
| **NYU** |  |  |  |  |  |  |
|  | non-NDD Pediatric | 0.80 | 0.17 | 0.91 | 0.06 | 23 |
|  | non-NDD Ped reduced (central only) | 0.84 | 0.14 | 0.91 | 0.07 | 23 |
|  | non-NDD Adult | 0.80 | 0.16 | 0.90 | 0.07 | 23 |
| **Overall** |  |  |  |  |  |  |
|  | non-NDD Pediatric | 0.82 | 0.11 | 0.91 | 0.05 | 135 |
|  | non-NDD Ped reduced (central only) | 0.80 | 0.10 | 0.91 | 0.05 | 135 |
|  | non-NDD Adult | 0.75 | 0.17 | 0.89 | 0.07 | 135 |
| **BCH** |  |  |  |  |  |  |
|  | NDD Pediatric | 0.76 | 0.16 | 0.89 | 0.07 | 41 |
|  | NDD Ped reduced (central only) | 0.71 | 0.17 | 0.87 | 0.09 | 41 |
|  | NDD Adult | 0.65 | 0.19 | 0.84 | 0.11 | 41 |
| **NIMH** |  |  |  |  |  |  |
|  | NDD Pediatric | 0.82 | 0.16 | 0.91 | 0.07 | 284 |
|  | NDD Ped reduced (central only) | 0.80 | 0.17 | 0.91 | 0.07 | 283 |
|  | NDD Adult | 0.78 | 0.21 | 0.89 | 0.08 | 284 |
| **NYU** |  |  |  |  |  |  |
|  | NDD Pediatric | 0.78 | 0.29 | 0.90 | 0.13 | 12 |
|  | NDD Ped reduced (central only) | 0.70 | 0.30 | 0.85 | 0.13 | 12 |
|  | NDD Adult | 0.68 | 0.26 | 0.84 | 0.12 | 12 |
| **Overall** |  |  |  |  |  |  |
|  | NDD Pediatric | 0.81 | 0.16 | 0.91 | 0.07 | 337 |
|  | NDD Ped reduced (central only) | 0.79 | 0.18 | 0.90 | 0.08 | 338 |
|  | NDD Adult | 0.75 | 0.21 | 0.89 | 0.09 | 337 |

**Table 3A:**
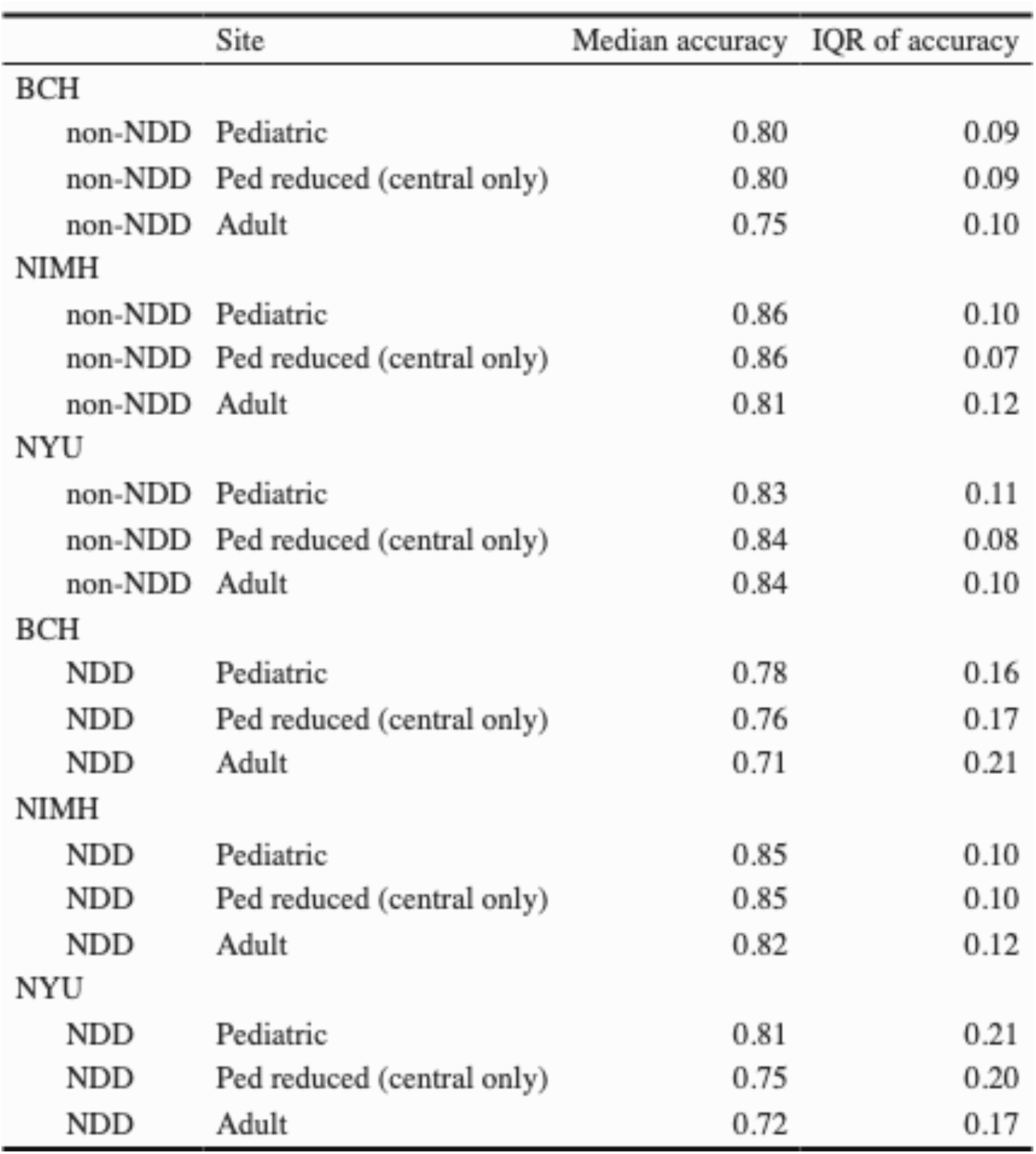
Transition accuracy between epochs flanked by similar epochs.

We observed similar results for epochs not adjacent to a transition (**Figure S1B; Table 3B**). Finally, we assessed whether automated staging performance changes in the context of NDD status, hypothesizing diminished performance of automated stagers in NDD cases. For all three stagers, we found significantly decreased performance in predicting sleep stage in NDD compared to typically developing cases (**Figure 1B**), but transition accuracy was not affected by NDD status (**Figure S1C**).

**Table 3B:** Transition accuracy between epochs at any type of transition.

| Site |  | Median accuracy | IQR of accuracy |
| --- | --- | --- | --- |
| BCH |  |  |  |
| non-NDD | Pediatric | 0.51 | 0.07 |
| non-NDD | Ped reduced (central only) | 0.50 | 0.07 |
| non-NDD | Adult | 0.47 | 0.06 |
| NIMH |  |  |  |
| non-NDD | Pediatric | 0.55 | 0.08 |
| non-NDD | Ped reduced (central only) | 0.56 | 0.10 |
| non-NDD | Adult | 0.53 | 0.08 |
| NYU |  |  |  |
| non-NDD | Pediatric | 0.54 | 0.14 |
| non-NDD | Ped reduced (central only) | 0.50 | 0.11 |
| non-NDD | Adult | 0.48 | 0.13 |
| BCH |  |  |  |
| NDD | Pediatric | 0.46 | 0.07 |
| NDD | Ped reduced (central only) | 0.43 | 0.08 |
| NDD | Adult | 0.41 | 0.08 |
| NIMH |  |  |  |
| NDD | Pediatric | 0.54 | 0.09 |
| NDD | Ped reduced (central only) | 0.54 | 0.09 |
| NDD | Adult | 0.52 | 0.09 |
| NYU |  |  |  |
| NDD | Pediatric | 0.49 | 0.12 |
| NDD | Ped reduced (central only) | 0.51 | 0.08 |
| NDD | Adult | 0.46 | 0.09 |

### Inferred properDes of sleep architecture are conserved between manual and automated staging

Rather than focus solely on epoch-wise concordance, we next asked whether automated stager-derived sleep features were comparable to those derived from manual staging. In RASP-M, we derived 1828 total features, comprising 61 macroarchitectural and NREM metrics seeded on either manual or automated staging. The metrics spanned six domains (macroarchitecture, spectral band power, spectral slope, slow oscillations, spindles, and interchannel connectivity, with metrics repeated for 10 EEG channels, seven band frequencies, and two spindle frequencies (slow and fast, 11 and 15 Hz, respectively; see Methods for details). Metrics derived from manual versus automated staging exhibited a high degree of correlation for all classes of features in the entire RASP-M cohort (mean *r* = 0.94, median *r* = 0.98; **Figure 2A; Table 4A**).

**Figure 2:**
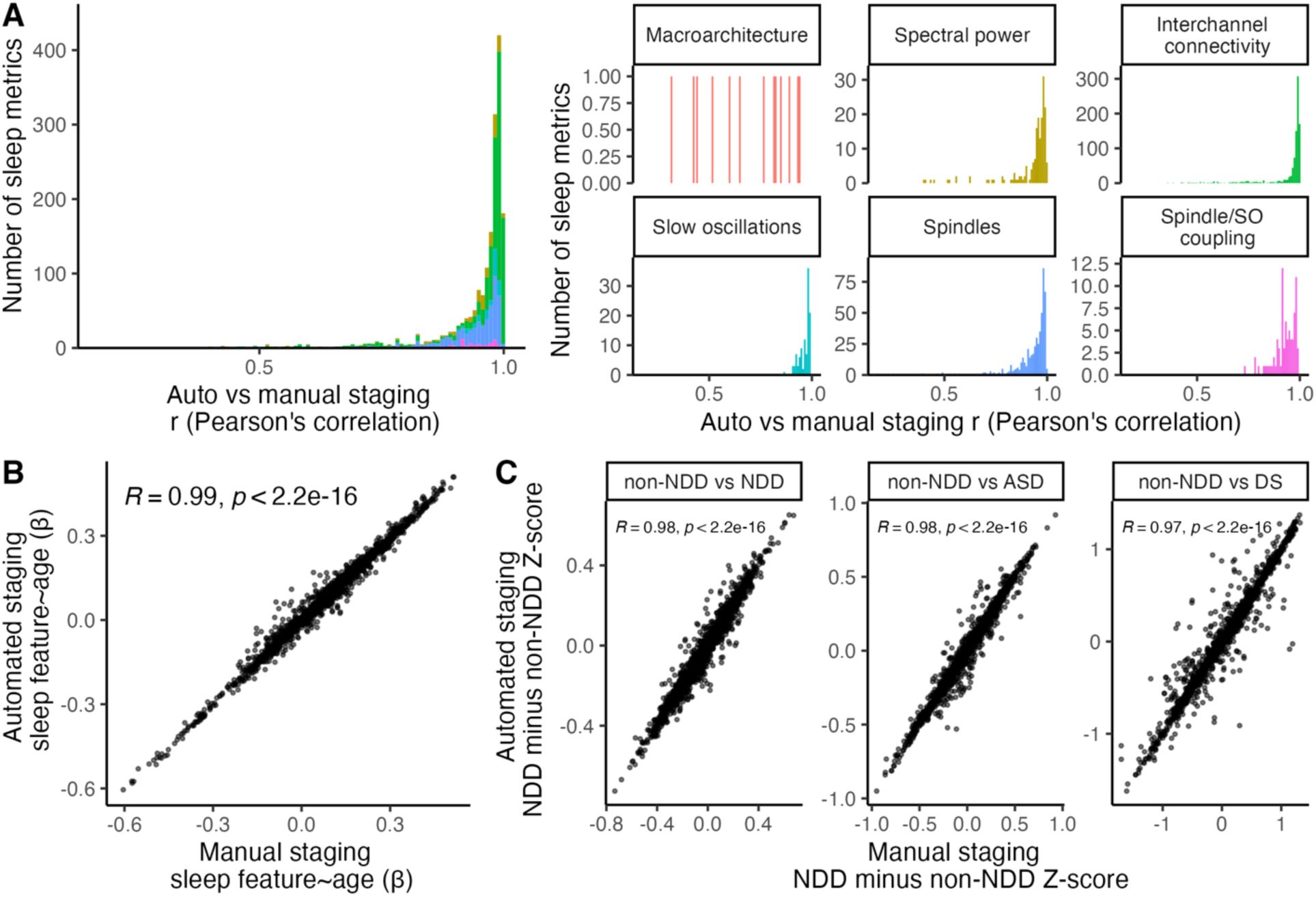
Automated staging conserves measures of sleep architecture including developmental trajectory and diagnosSc characterisScs. A) Distribution of Pearson’s correlation coefficient comparing automated versus manual staging across all studied sleep metrics (lev), and by sleep feature class (right). B) Correlation of developmental changes in sleep features derived from manual versus automated staging. C) Correlation of differences in sleep architecture between NDD minus non-NDD across all NDDs (lev), ASD (middle), and DS (right). For both B and C, each point represents a sleep metric. X and Y axes denote the beta of the age and diagnosis coefficient, respectively, in the regression equation. R denotes Pearson’s correlation coefficient between betas and associated p-value is shown.

**Table 4A:** Correlation of sleep features derived from manual vs automated staging in RASP subjects ≥ 2.5 years.

| Class | Median | IQR |
| --- | --- | --- |
| Macroarchitecture | 0.76 | 0.33 |
| Spectral power | 0.96 | 0.05 |
| Interchannel connectivity | 0.99 | 0.02 |
| Slow oscillations | 0.98 | 0.04 |
| Spindles | 0.96 | 0.07 |
| Spindle/SO coupling | 0.93 | 0.07 |

However, correlations between macro-architectural metrics from manual and automated staging were lower (mean *r* = 0.69, median *r* = 0.76; **Figure 2A, Table 4A**). On closer inspection, the proportion of lme spent in individual NREM stages has the lowest correlation across macroarchitecture features (**Table 4B**), paralleling previously reported uncertainty in manual staging of NREM stages [45–49].

**Table 4B:** Correlation of macroarchitecture features derived from manual vs automated staging in RASP subjects ≥ 25 years.

| Variable | r |
| --- | --- |
| Sleep period time (onset to wake) | 0.93 |
| Total sleep time | 0.94 |
| Total wake time | 0.89 |
| Wake time after sleep onset | 0.82 |
| Sleep onset latency | 0.85 |
| % NREM | 0.51 |
| % N1 | 0.44 |
| % N2 | 0.31 |
| % N3 | 0.42 |
| REM onset latency | 0.60 |
| Sleep efficiency | 0.83 |
| Sleep maintenance efficiency | 0.76 |
| Sleep fragmentation index | 0.65 |

We next assessed whether analyses based on automated staging recapitulated developmental and diagnostic changes in sleep architecture. In RASP-M, we regressed individual sleep features on age, correcting for sex, study site, and diagnosis. There was a high degree of correlation (r = 0.99) between developmental changes in sleep features derived from manual and automated staging, as based on the beta coefficients for age from the two sets of analyses (**Figure 2B**), which held across all classes of feature (**Figure S2A**). Second, we asked whether analyses based on automated staging accurately reflected changes in sleep macro- and microarchitecture in NDDs, focusing on DS and ASD. For each sleep metric, we derived the mean group differences between non-NDD and NDD, ASD, or DS cases, correcting for the variables described above. Again, we observed a high degree of correlation (r =0.98 for non-NDD versus NDD, ASD, or DS cases) between results based on manual and automated staging (**Figure 2C**) for all feature classes (**Figure S2B**). Taken together, these results directly demonstrate that our pediatric stager-derived inferences on developmental and diagnosis-dependent changes in sleep architecture recapitulate those derived from manual staging. This supports the application of the pediatric stager to all RASP individuals as a basis of uniform analyses across sites. All results that follow are based on the full pediatric model staging.

### Conserved developmental changes in sleep architecture in RASP cohorts

Next, we focused our analysis on data from children without NDDs aged 2.5 years and older from all five RASP cohorts (**Figure 3A)**. Multiple metrics showed age-dependent changes consistent with prior literature. Total sleep time decreased with age whereas sleep fragmentation increased, reflected by decreased sleep maintenance efficiency, increased sleep fragmentation index, and increased wake lme aver sleep onset **(Figure 3B**, **Table 5).** Across age, the proportion of lme spent in N2 sleep increased while that spent in N3 decreased (**Figure 3C**; **Table 5).** Microarchitectural metrics such as NREM band power, sleep spindle features, and slow oscillation measures also exhibited marked developmental changes (**Table 6**). In the RASP cohorts, absolute slow and delta band power decreased across development, while sigma band power increased (**Figure 4A**; **Table 6)**. Fast and slow spindle density increased across development, and intra-spindle frequency change (chirp), decreased across development (**Figure 4B**; **Table 6**). Slow oscillation duration, rate, and negative amplitude increased with age, while peak-to-peak amplitude decreased (**Figure 4C**; **Table 6**). These findings recapitulated known developmental changes in sleep architecture [28]. Thus, developmental changes in sleep architecture are conserved in the RASP cohorts.

**Table 5:** Developmental changes in sleep macroarehitecturc features in RASP non-NDD ≥ 2.5 years.

| Metric | $\beta$ | P-value | |
| --- | --- | --- | --- |
| **Sleep time and fragmentation** |  |  |  |
| Total sleep time | -0.100 | 0.0075 | ** |
| Sleep maintenance efficiency | -0.140 | 0.00017 | *** |
| Sleep fragmentation index | 0.300 | 6.4e-16 | *** |
| Wake after sleep onset | 0.140 | 0.00019 | *** |
| **Proportion of time in sleep stage** |  |  |  |
| N1 | 0.050 | 0.19 | ns |
| N2 | 0.150 | 6e-05 | *** |
| N3 | -0.190 | 2.9e-07 | *** |
| R | 0.019 | 0.61 | ns |

**Tabic 6:**
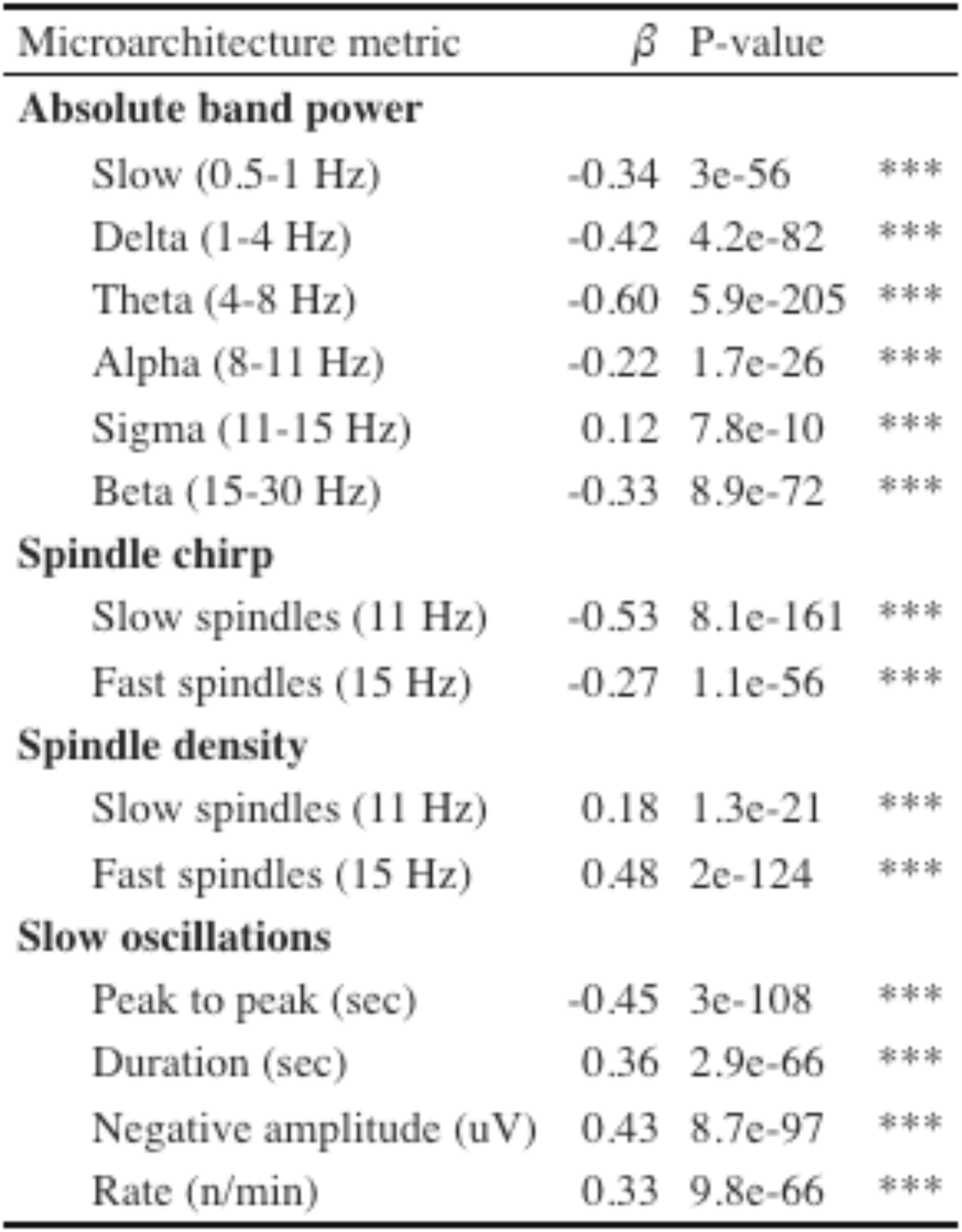
Developmental changes in sleep microarchitectural features in RASP non NDD.

**Figure 3:**
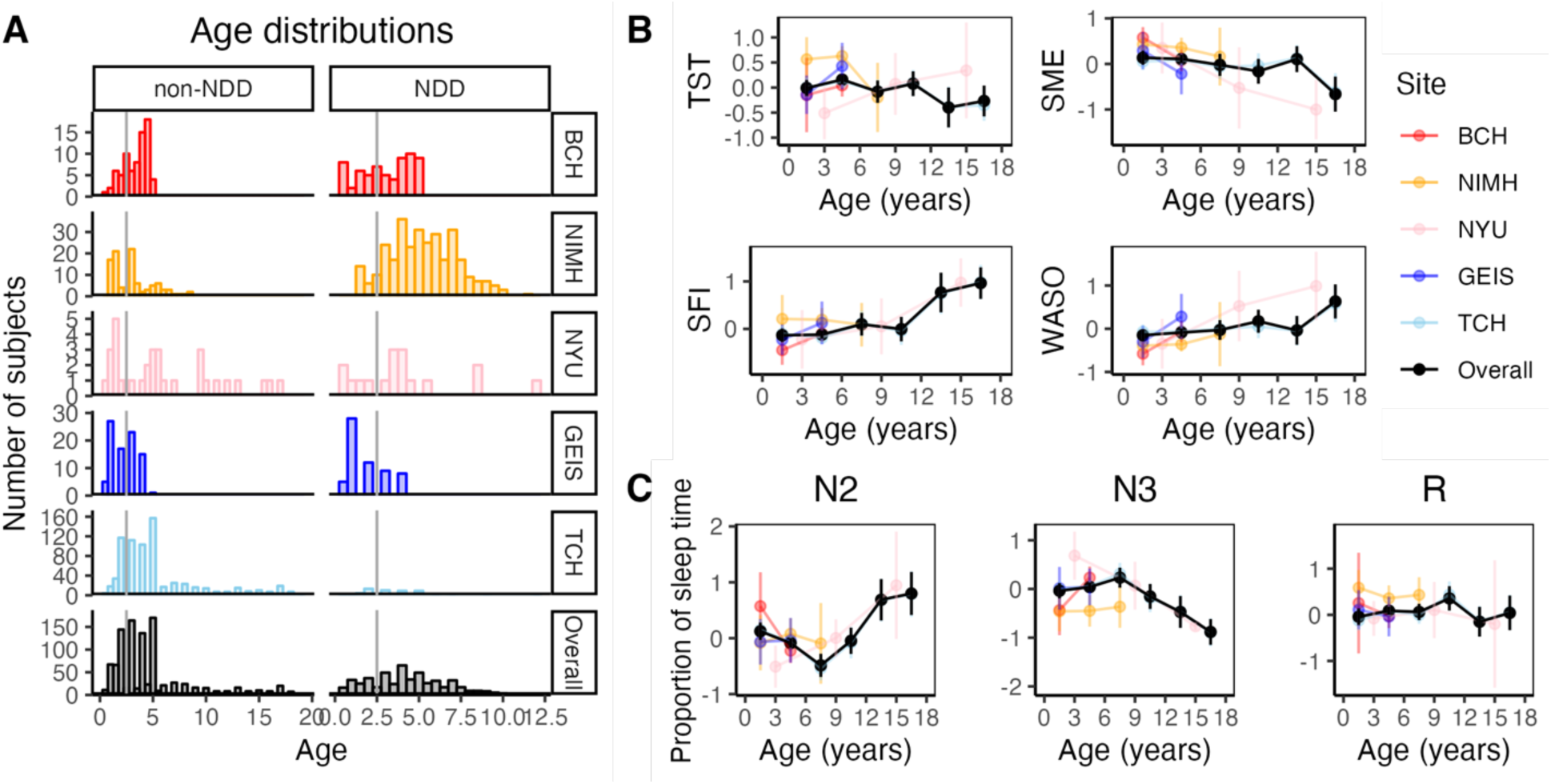
The RASP cohorts demonstrate known developmental changes in sleep macroarchitecture. A) RASP age distributions by study site. All datasets underwent automated staging with a pediatric stager to extract sleep architectural metrics. B) Total sleep lme (TST), sleep maintenance efficiency (SME), sleep fragmentation index (SFI), and wake aver sleep onset (WASO) in the RASP cohorts across development. C) Proportion of sleep lme spent in N2, N3, and R stages across development. All sleep metrics in this figure and all subsequent figures are reported as Z-scores across individual sites. The overall trace (black) represents a compilation of Z-scores.

**Figure 4:**
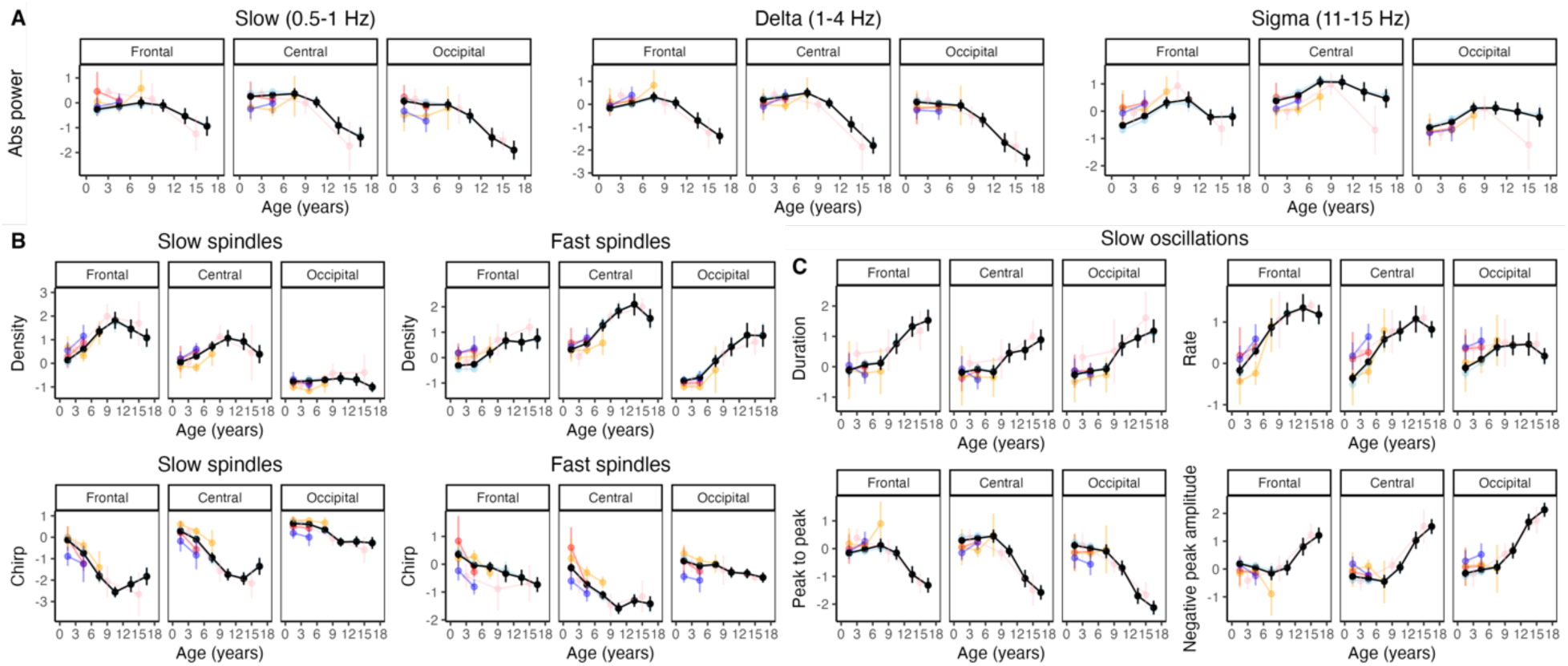
Developmental changes in NREM microarchitectural metrics are conserved in RASP. A) Absolute power of slow, delta, and sigma band frequencies, B) slow and fast spindle density (top) and chirp (bottom), and C) slow oscillation duration, rate, peak-to-peak amplitude, and negative peak amplitude (lev to right, top to bottom) across age.

### Consistent differences in sleep architecture across cohorts in DS but not ASD

Using the RASP cohorts, we next examined how sleep architecture differs in the context of DS. We examined 1351 total sleep features, representing 79 metrics in the six domains studied above (macroarchitecture, spectral band power, spindles, slow oscillations (SO), spindle/SO coupling, and interchannel connectivity). These were repeated for paired and averaged frontal, central, and occipital channels, seven band frequencies, and three sleep stages (N2, N3, and REM). In the RASP dataset, we found 66 metrics spanning all 6 domains that differed between non-NDD and DS cases (using FDR-adjusted p-value < 0.05; **Fig S3A**). To assess whether sleep architecture differences were consistent across independent cohorts, we used DS cases from the NCH-SDB. We found 73 metrics across all six domains that were different compared to non-NDD NCH-SDB cases (**Fig S3C)**. RASP and NCH-SDB DS cases shared 55 metrics across all six domains that deviated from their respective non-NDD cohorts (total 317 features when accounting for all channels, band frequencies, and sleep stages; i.e., 85% of the RASP cohorts features were conserved in a separate cohort). Consistent with this, there was a significant degree of correlation between RASP and NCH-SDB DS vs non-NDD sleep metric changes (r = 0.78 in the beta coefficients of the diagnosis covariate across all metrics; **Fig 5A; Fig S3E**). Thus, DS-associated sleep architecture changes were broadly consistent across independent cohorts.

**Figure 5:**
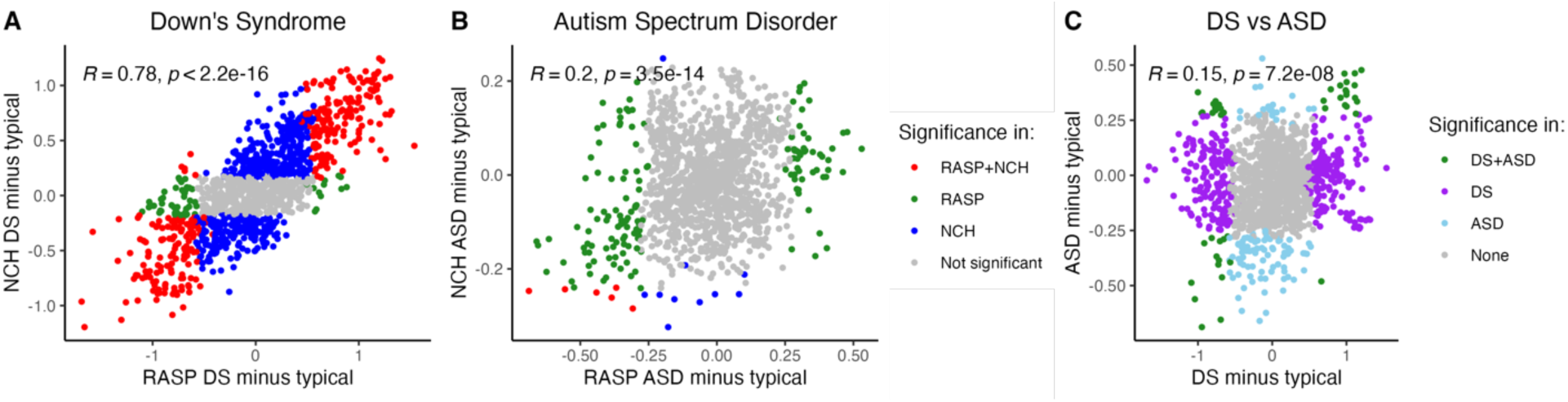
Sleep architecture changes related to NDD diagnosis are consistent across RASP and NCH-SDB cohorts in DS but not ASD. A) Cross-cohort comparison of sleep architecture metrics in DS vs typical development. Each point represents an individual sleep feature. Red denotes features that are significant across both RASP and NCH, green are features significant in only RASP, and blue are features significant in only NCH. B) Cross-cohort comparison of sleep architecture metrics in ASD vs typical development. C) Cross-diagnosis comparison of sleep architecture changes in DS vs ASD. Pearson’s correlation coefficient is shown for all figures.

Next, we asked how sleep architecture differs in the setting of ASD. In the RASP cohorts, 42 metrics across all domains except spindle/slow oscillation coupling were different between non-NDD and ASD cases (**Fig S3B)**. These features did not show a high degree of correlation or overlap between DS and ASD; only 15 significantly different sleep metrics (32 total features) were shared between DS and ASD (**Fig 5C**), suggesting sleep architecture changes in DS may be specific to DS. In NCH-SDB ASD cases, 10 sleep metrics across four domains (macroarchitecture, spectral band power, slow oscillation, and interchannel connectivity) deviated from non-NDD cases (**Fig S3D**), and only 4 metrics (from spectral band power and interchannel connectivity) were shared between RASP and NCH-SDB (**Fig 5B; Fig S3F)**. Further, differences in sleep architecture in ASD were only modestly correlated across RASP and NCH-SDB (r = 0.21) (**Fig 5B**).

### PredicDng brain age based on pediatric sleep EEGs

To summarize and compare the differences in sleep architecture features we observed in DS and ASD, we constructed a model to predict brain age in the RASP NDD cohort. Using pediatric cases in the NCH-SDB, PATS, and CHAT, we fit a linear regression of sleep measurements on age (correcting for sex, race, and study site) to identify relevant features to include in our final model. As microarchitecture features alone predicted age as well as models additionally including macroarchitectural measures [28], we restricted feature identification to sleep microarchitectural metrics. Out of 1338 features, we identified 448 features for which age explained at least 2.5% of the variance in sleep feature (**Figure 6A**). As an example, REM relative alpha band power but not N2 theta band power kurtosis measured from occipital EEG channels exhibited a robust correlation with age (**Figure 6B**). Given the goal of building a robust model transferable between cohorts, we excluded metrics with significant inter-site variability at this stage: for example, although spectral slope measures (estimated by fixng log-log linear regression models on absolute power in the 30-45 Hz frequency range) were significantly associated with age, we noted these effects were not consistent across all three studies (likely driven by study-dependent differences in filtering higher frequency activity prior to exporting recordings) and thus excluded these metrics (**Figure 6B**). Of note, removing race as a covariate did not affect the correlations between age and individual sleep metrics, with high correlation between FDR-adjusted p-values when including and excluding race (r = 0.99).

**Figure 6:**
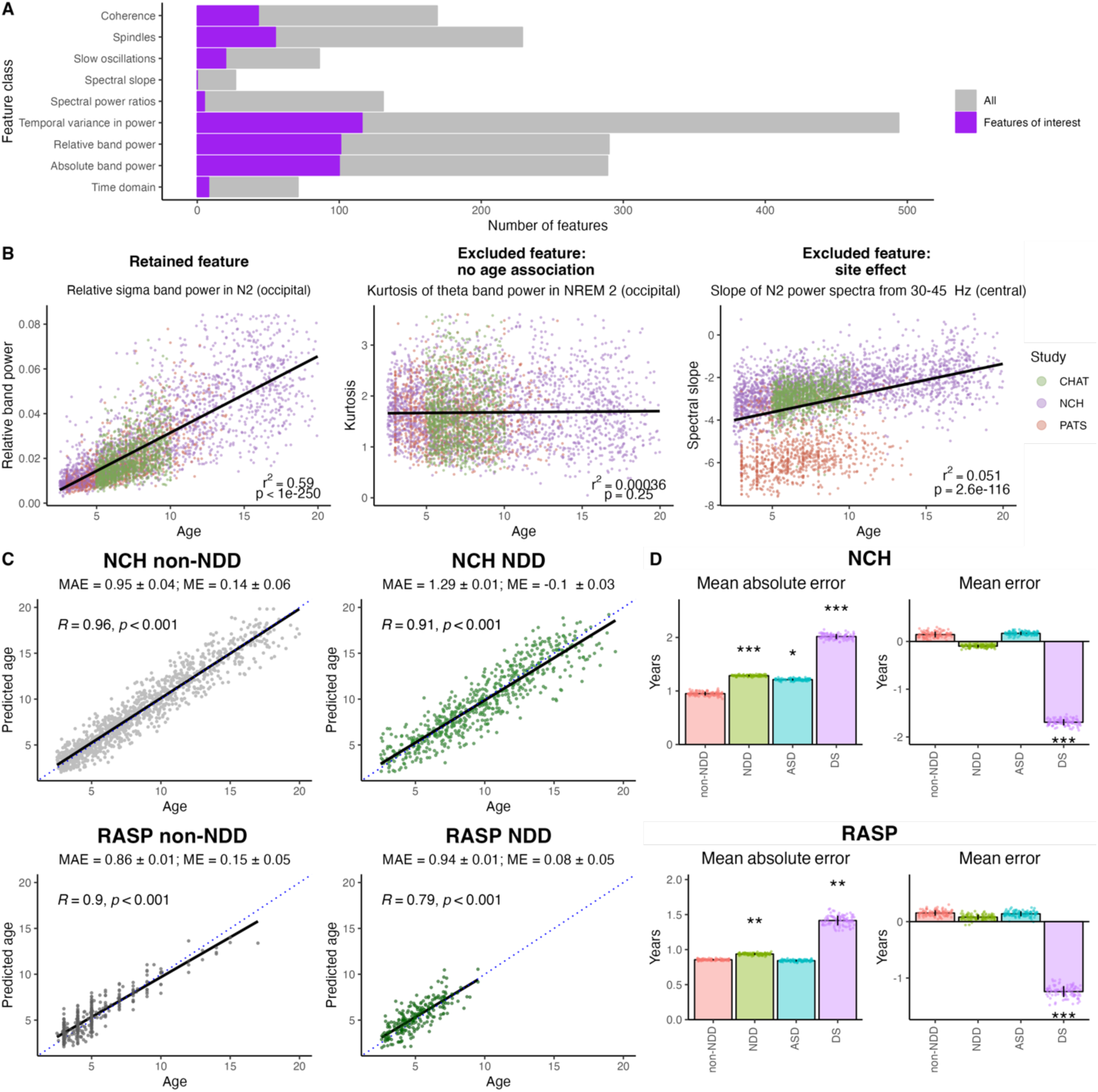
Sleep feature selecSon and model performance on NCH holdout, RASP non-NDD, and NDD cases recapitulates younger predicted brain age in DS. A) Sleep features of interest (purple) compared to all features (gray), broken down by feature class. Sleep EEG features highly correlated with age (purple) in non-NDD CHAT, NCH, and PATS cohorts were identified as initial features of interest for building a model for predicting age. B) Example developmental changes in three different sleep metrics. Age predicts a large proportion (63%) of the variation in REM relative alpha band power (lev) but predicts close to none of the variation in NREM theta band power kurtosis (middle). Spectral slope (log-log linear regression of the power spectra from 30-45 Hz) in N2 versus age demonstrates a metric with significant intersite variability. P-values indicate significance of correlation between age and sleep metric, correcting for sex, race, and study site. C) Measured vs predicted age by final model in NCH and RASP non-NDD vs NDD cohorts. Mean absolute error (MAE) and mean error (ME) values ± standard deviation across were derived from resampling the training set and testing the final model on each group 100x; NCH non-NDD was resampled as 30% of the NCH training dataset for each run. Pearson’s correlation (R) is reported for a single run for each group. D) MAE and ME for NDD groups in comparison to respective non-NDD groups across 100x resampling of the training dataset. Bars represent average error across all 100 samplings, and error bars represent standard deviations. Median p-values are reported for comparisons within samplings, comparing each group to non-NDD, correcting for age and sex in addition to study site in RASP (* p<0.05, ** p < 0.01, *** p < 0.001).

We first sought to assess how well our model predicted age in typically developing individuals. We trained our model on individuals without an NDD diagnosis aged 2.5 to 18 years from NCH-SDB, CHAT, and PATS and predicted age in two separate cohorts: 1) a hold-out 30% of the non-NDD NCH-SDB group and 2) the RASP cohorts. To assess error in age predictions, we performed repeated hold-out testing to estimate model performance across 100 repetitions, resampling the NCH-SDB holdout and training set during each repeat. In the NCH-SDB holdout set, the mean absolute error (MAE) was 0.95 years (95% CI of 0.06 years). In the non-NDD RASP cohorts, MAE was 0.81 years (95% CI of 0.01 years). Pearson’s correlation coefficients between actual and predicted age in the NCH-SDB holdout set and RASP non-NDD cohorts were 0.96 and 0.9, respectively **(Figure 6C**). Altogether, these results suggest our model predicts chronologic age with a high degree of accuracy in children without an NDD diagnosis across independent cohorts.

How does NDD diagnosis affect brain age prediction? Compared to non-NDD cases, there was decreased correlation between actual and predicted age (**Figure 6C**) as well as increased MAE (**Figure 6D**). In DS, our model predicted increased MAE and younger brain age in both NCH-SDB and RASP cohorts (**Figure 6D**), recapitulating previous findings [28]. NCH-SDB ASD cases had increased MAE compared to non-NDD controls, which was not observed in RASP cases. Predicted age difference was not significantly different in either of the ASD cohorts compared to non-NDD individuals (**Figure 6D**). Thus, we demonstrate in an independent cohort that individuals with DS have a younger predicted brain age compared to non-NDD cases.

Although several sleep metrics were significantly different between ASD and typically developing cases, we did not find a difference in predicted brain age in RASP ASD cases, perhaps reflecting the heterogeneity seen across ASD. We predicted there would be increased variance in sleep metrics in ASD compared to typical development. To account differences across study sites, we first regressed site on individual sleep metrics, correcting for age, sex, and diagnostic status, to obtain site-adjusted residuals. We compared the variances of these residuals in age- and sex-matched non-NDD cases to that of either DS or ASD cases. In DS, 15 metrics had significantly different variances. Of these, 10 had a higher variance in the DS group. In ASD, of the 297 metrics with significantly different variances, 281 metrics (95%) had higher variance in ASD cases (**Figure S4**). Overall, significant differences were broadly distributed with no obvious patterning across sleep metric class, sleep stage, frequency band, or EEG channel.

### Feature selecDon affects the generalizability of model predicDons

Feature selection is a crucial step for optimizing model performance and efficiency while offering valuable insights into processes underlying the data [50]. Our initial model considered features across all available channels and sleep stages that exhibited changes with age. In a series of secondary analyses, we asked how sleep feature selection affected age prediction. For example, do estimates of predicted age differences (PADs) based on NREM sleep provide similar information those based on REM, and is one class of predictor more sensitive to the changes observed in DS?

We generated sleep stage-specific models by restricting to features from only N2, N3, or REM. To account for differences in feature number when comparing stage-specific models with our full model, we included a fourth model with an equivalent number of randomly selected features from all sleep stages (matching the average number of features across N2, N3, and REM-specific models). Using repeated k-fold cross validation, our original model gave the lowest MAE and highest coefficient of determination (R^2^) compared the four other models. Using only features from REM resulted in the worst performing model (reducing R^2^ = 0.88 to R^2^ = 0.73 ) (**Figure 7A**). Next, we asked how stage-specific models affect the age prediction of non-NDD versus NDD cases. For DS in NCH-SDB, all four models predicted a significantly younger age gap in DS compared to non-NDD cases (**Figure 7B**). This effect was attenuated in the N3 and REM models for NCH-SBD DS cases. In contrast, when we applied these four models to RASP DS cases, N3 and REM feature models did not result in any difference in predicted age in non-NDD versus DS **(Figure 7B**). Age predictions on NCH-SDB ASD cases exhibited more variability across models, with older predicted age by REM and random subset feature models (**Figure 7B**). These effects were not consistent in RASP ASD cases, which did not show any differences in predicted age under any of the models **(Figure 7B**). Thus, subsexng features by sleep stage decreases model performance and generalizability of age prediction across cohorts, suggesting that all stages provide some degree of independent information relevant to typical and atypical development.

**Figure 7:**
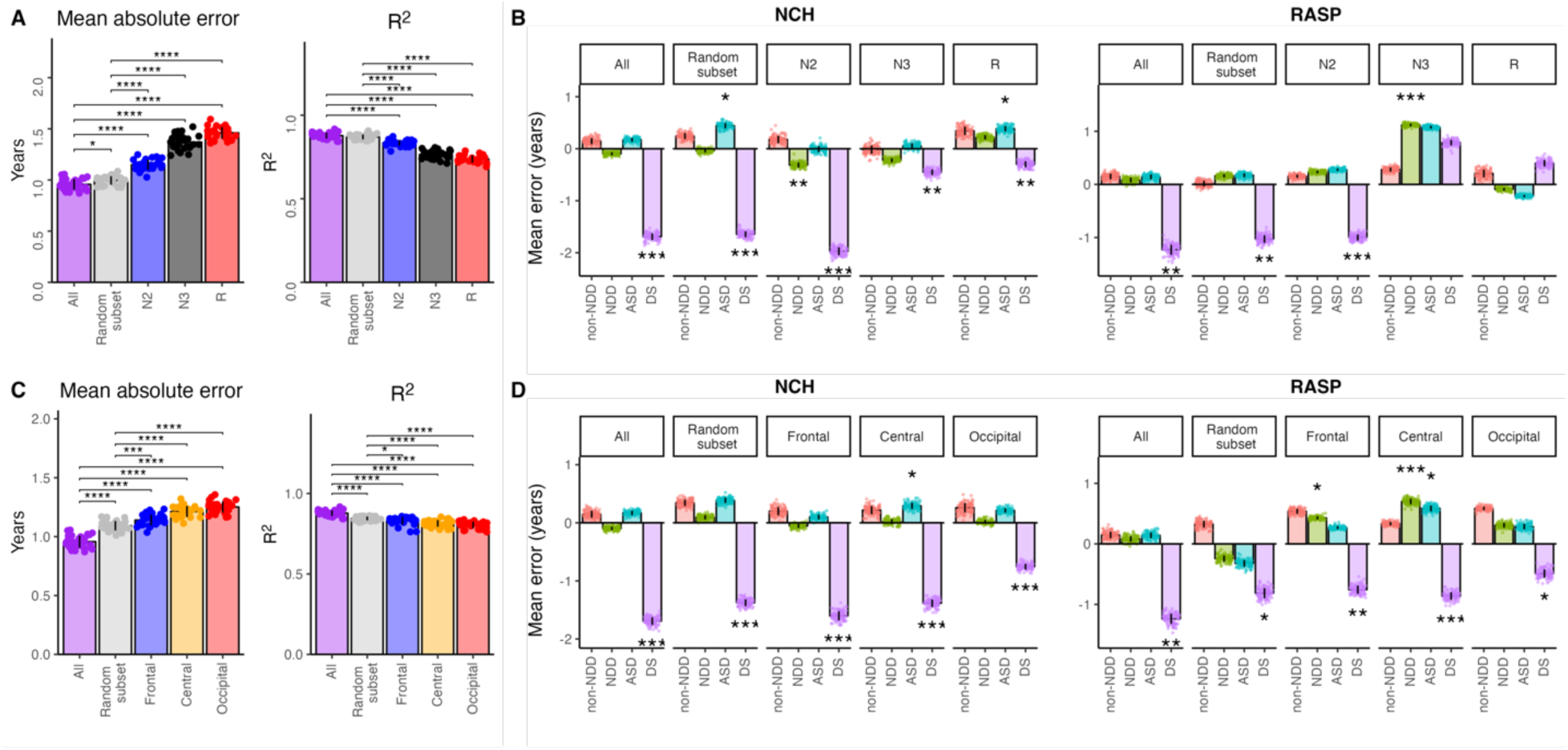
Feature selecSon changes age predicSon model performance. Model performance based on sets of sleep features from specific A) sleep stages or C) EEG channels. Performance is reported as mean absolute error and R^2^ (variance explained) across all folds of repeated k-fold validation (3 repeats and 10-fold) and compared to general predictive model (purple columns; multiple T-tests with Bonferroni correction). Age gap (i.e., mean error) across groups according to predicted age from sleep features from specific B) sleep stages or C) EEG channels across 100x resampling of the training dataset. Bars represent average error across all 100 samplings, and error bars represent standard deviations. Median p-values are reported for comparisons within samplings, comparing each group to non-NDD, correcting for age and sex in addition to study site in RASP (* p<0.05, ** p < 0.01, *** p < 0.001).

Can we achieve similar conclusions with fewer EEG channels? Although PSGs are the gold standard, reduced montages and single channel devices would allow for more accessible, scalable home-based longitudinal recording. To test whether brain age prediction is consistent across models with features from fewer EEG channels, we repeated our analysis with features from specific channels. Again, we included a random subset of an equivalent number of features (again, matching the average of the number of features from each individual model) here from single EEG channels (i.e. excluding cross-channel coherence metrics). As observed in the stage-specific models, our original multi-channel model had the best performance (lowest MAE and highest R^2^) compared to the models with restricted features (**Figure 7C**). In both NCH-SDB and RASP, effects on DS and ASD cohorts were more uniform, apart from an older predicted age in ASD with the central channel feature model **(Figure 7D)**. Together, these findings suggest including information from all sleep stages improves the applicability of our model, and a reduced number of EEG channels results in worse model performance but is sufficient for broad comparisons of brain age across diagnostic status.

## Discussion

Here, we introduced a collection of clinical pediatric overnight polysomnography recordings collected retrospectively across five different institutions, aiming to support pediatric sleep research including studies on brain development. Disruptions in specific aspects of sleep architecture provide a unique window into affected neural circuitry. Thus, abnormalities in early life sleep architecture represent a readout for neural circuit dysfunction and abnormal brain development. Here, we recapitulated our previous findings that sleep EEG in children with DS predicts younger brain age [28]. This consistency across independent cohorts points to robust and conserved alterations in DS sleep architecture. Indeed, at the level of individual sleep metrics, we demonstrate consistency in changes in sleep architecture changes across DS cases in the RASP cohorts and NCH-SDB. Further understanding the range of sleep problems in individuals with NDDs and how these relate to abnormal brain development necessitates large datasets to define sleep EEG-based diagnostic groups within and across NDDs. Linked with longitudinal and genetic data, future studies of sleep abnormalities across patients with neurodevelopmental disorders could classify biomarkers of diagnostic groups and reveal mechanistic links to circuit-level disruptions. Our work points to the feasibility of using retrospective clinical data to this end.

Given the retrospective nature of our dataset and analyses, several caveats should be noted. First, obstructive sleep apnea (OSA) is highly prevalent in Down’s syndrome [51], presenting a confounding variable that might affect brain function and estimates of brain age. Children with severe OSA may have decreased deep (N3) sleep, reduced deep sleep efficiency, and increased sleep latency compared to healthy controls [52]. Sleep microarchitecture measures are also affected in OSA; for example, there is a reduction in slow wave sleep in the context of OSA as well as changes in sleep spindle measurements [53,54]. Reassuringly, inclusion of OSA severity (apnea hypopnea index) as a covariate in sleep EEG-based age prediction models in our previous analysis did not affect brain age prediction [28]. Second, we did not account for heterogeneity within NDD groups in our present analyses. Indeed, we found poor correlation between sleep metrics from RASP and NCH-SDB ASD cases, suggestive of differences between ASD cases across these two cohorts. In addition, we observed increased variance in individual sleep metrics from the RASP ASD cohort. Together, our results suggest sleep changes in the context of ASD are heterogeneous, paralleling known variability in symptomatology. Future studies investigating whether sleep alterations in specific NDDs cluster by severity of behavioral presentation would better elucidate whether specific sleep disruptions are associated with functional status. Finally, medication information was not available for the RASP cohorts, presenting another variable that might affect sleep.

The prevalence of sleep disturbances across NDDs may reflect similarities in underlying aberrant sleep physiology across these disorders. Sleep in early life is critical for brain development. Does disrupted sleep in NDDs impair neurodevelopment? Sleep problems are strongly associated with cognitive and behavioral deficits in both typically developing children and those with NDDs [55–57]. In individuals who later go on to develop ASD or DD, sleep problems per caregiver report in early childhood are associated with more ritualistic and compulsive behaviors later in childhood [58]. Additionally, caregiver-reported sleep onset problems in infants are associated with alterations in developmental trajectory of hippocampal volumes across childhood [59]. Animal models of NDDs suggest sleep disruptions arise early in life and are linked to abnormal development of sleep-associated neural circuits [60–62]. In mice with mutations in SHANK3, an ASD-associated gene, longitudinal EEG recordings reveal decreased sleep, disrupted sleep architecture, and abnormal sleep homeostatic responses emerge during early development [62]. Together, these findings suggest sleep disruptions may be a core feature of NDDs that arise from abnormal development. However, whether sleep-based interventions in early life result in less severe behavioral and/or cognitive symptoms later in life is unclear. This remains a crucial unanswered question regarding whether NDD-related sleep disturbances drive abnormal neurodevelopment. Additionally, most clinical studies include caregiver reported metrics of sleep, while objective measures of sleep architecture from EEG recordings are lacking. Correlating patient sleep architecture disruptions with specific abnormalities in animal models of NDDs to uncover conserved neurocircuits would greatly aid the assessment and development of sleep-targeted interventions, which are currently limited in pediatric populations [63].

To facilitate our use of retrospective clinical datasets, we first faced the challenge of missing stage annotations across several cohorts, combined with the likelihood that generic stagers trained on primarily adult data may not perform optimally. We therefore developed a pediatric stager to support all subsequent analyses of these recordings. As well as high epoch-wise kappa statistics, we show that automated staging is effectively comparable to manual staging in terms of the statistical inferences made on the sample-level. Our results support previous reports [33] that stagers derived from pediatric sleep data are better suited for automated sleep staging in pediatric cohorts compared to existing adult stagers. Of note, macro-architecture metrics appeared more sensitive to differences between manual versus automated staging compared to microarchitectural feature classes (**Figure 2A, Table 4A-B**).

Proportion of lme spent in individual NREM stages had the lowest correlation between manual and automated staging, although this arguably reflects the known lower inter-rater reliability in distinguishing NREM stages [45–49]. Reassuringly, wake and sleep durations derived from automated staging correlated well with those from manual staging (**Table 4B**).

PSG is currently the gold standard for sleep studies but is resource- and time-intensive, making large scale data collection difficult. Automated classification of sleep stages using our pediatric stager achieves a high level of agreement with manual staging, outperforms existing adult stagers, and conserves sleep architecture metrics. Importantly, consistent automation is promising for overcoming bottlenecks inherent in the manual interpretation of sleep studies, especially in the context of assembling archived clinical studies from different sources. The technical aspects of PSG present another limitation: PSGs are performed in specialized lab sexngs, which may not be tolerated by younger children. Although both automated staging and EEG-based age prediction benefitted from an increased number of channels, it is worth noting that single channel models were sufficient for both staging and diagnostic comparisons of age prediction, respectively. This is significant, as it suggests that ambulatory EEG devices with limited montages including “wearables” can provide a scalable framework for biological age prediction. Specifically, we found predicting age using features from fewer EEG channel measurements resulted in consistent age predictions across diagnostic groups. A reduced number of EEG channels may be sufficient for drawing conclusions about brain health and function in the context of sleep. Developing home-based recording techniques with a limited number of EEG channels for patients with NDDs would increase the accessibility of studying sleep in this population. Coupled with automated staging analysis, such an approach would allow for rapid data collection and analysis. enhance performance and How might feature selection in brain age prediction models deepen our understanding of mechanisms underlying NDDs? Sleep EEG in children with DS predicts younger age in multiple cohorts; however, in both structural brain MRI studies [64] as well as DNA methylation measures from brain and blood, adults with DS have older predicted age [65]. While these findings suggest abnormal aging in later life, these effects are not restricted to adult patients: DNA methylation-based “epigenetic clock” in blood samples from neonates with DS also pointed towards accelerated aging in very early life [66]. These disparate conclusions could be based in varied biological insights provided by different methodologies. Sleep EEG-based age predictions stem from functional differences in the sexng of NDDs. For example, children with DS have delays in psychomotor development reflected by significantly younger developmental age compared to typically developing children [67]. These findings parallel our findings that children with DS have younger predicted age based on sleep architecture. Of note, age prediction using features from specific sleep stages diminishes model performance and the ability to generalize findings across cohorts (**Figure 7**). This may reflect how distinct facets of sleep architecture are affected in the context of different NDDs [23], as well as how specific sleep stages may inform different aspects of brain function. Indeed, sleep features derived from NREM versus REM stages in mobile sleep EEG recordings resulted in differences in predicted brain age [68]. Multimodal approaches integrating imaging, genomics, and functional methods such as sleep EEG may improve age prediction by providing more information about the mechanisms underlying biological differences. Analogously, incorporating different structural imaging modalities in a brain age prediction model improves brain age prediction [64].

What insights does the predicted age metric provide regarding neurocircuit mechanism of disease in NDDs? Predicted brain age based on structural and functional MRI metrics in 504 participants aged 36-100 years explained only an additional 1.6% of the variation in fluid cognition in healthy adult subjects [69]. Using our age prediction model, NCH-SDB but not RASP ASD cases exhibited increased mean absolute error between predicted and measured age, but no difference in predicted age gap (**Figure 6D**). Furthermore, RASP ASD cases had differences in specific sleep metrics compared to non-NDD cases, but these were not reflected in NCH-SDB ASD cases. Compared to typically developing cases, RASP ASD cases additionally exhibited increased variance across individual metrics. While these results likely reflect the heterogeneity in ASD clinical presentation, predicted brain age may also not adequately capture the relevant differences in ASD neurophysiology. Focusing on other outcomes (i.e., cognition or behavioral symptoms) may yield more insight into how sleep architecture disruptions reflect specific circuit dysfunction. Future studies comparing cross-modal changes between typically developing and NDD patients as well as relating overnight sleep architecture to additional measures besides brain age would be helpful for clarification of how sleep disturbances reflect disruptions in brain function.

## Methods

### ParDcipants

We used overnight PSGs from five institutions (the RASP cohorts). The NIMH cohort consisted of 212 unique individuals (186 over the age of 2.5 years), with repeat longitudinal encounters for a total of 413 recordings (345 over the age of 2.5 years). We treated each recording individually for subsequent analyses. We additionally used overnight PSG data from three other pediatric samples: 1) the Nationwide Children’s Hospital Sleep Databank (NCH-SDB) [36,37], the largest pediatric sleep clinic cohort of 3,984 sleep studies on 3,673 unique pediatric patients; 2) the Child Adenotonsillectomy Trial (CHAT) [38], which is comprised of 1,244 pediatric patients aged 5 to 9 from six US clinical centers screened for OSA in consideration of testing adenotonsillectomy for treatment of OSA; and 3) the Pediatric Adenotonsillectomy Trial for Snoring (PATS) [39,40], which included 1008 sleep studies from 704 children ages 3 to 13 years from six US clinical centers screened for snoring in consideration of testing adenotonsillectomy for treatment of primary snoring. NCH-SDB, CHAT, and PATS data were available through the National Sleep Research Resource (NSRR) [36].

### Automated sleep staging

We developed and evaluated an automated stager (POPS) implemented within the Luna toolset, using three models: a) a single (central) channel model trained on adult data, b) a single (central) model trained on pediatric data, and c) a multi-channel (frontal, central and occipital EEG and EOG) model based on pediatric data.

#### Training and test data

The adult model was trained on 3276 individuals selected from six NSRR cohorts (CHAT, CCSHS, CFS, MESA, MrOS and SHHS) with 600 held out for testing the trained model. The pediatric models were trained on 3582 individuals selected from two pediatric cohorts (NCH-SDB and CHAT) with 580 held out for testing the trained model. Single-channel models were trained using both C3-M2 and C4-M1 central EEGs, with each record entered independently into the training model as we assumed that metrics derived from each channel would exhibit effectively identical stage-dependent associations. Likewise, multi-channel models were in fact based on three EEG channels, either lev hemisphere (F3-M2, C3-M2 and O1-M2) or right hemisphere (F4-M1, C4-M1 and O2-M1) channels, as well as a single EOG. For all models, when used to predict sleep stages in new data, we made two initial sets of predictions, based on either the lev or right channel(s). For each epoch, the prediction consists of five posterior probabilities that sum to 1.0: P(stage|data) for N1, N2, N3, REM and wake. We then combined lev and right predictions by selecting for each 30 second epoch the prediction (either lev or right) with the highest confidence, where confidence was defined as the maximum posterior probability.

#### Per-epoch model features

Each EEG channel was resampled to 128 Hz, scaled to micro-volt units and band-pass filtered using a zero-phase Kaiser window FIR with transition frequencies at 0.3 and 35 Hz. We computed a time-domain normalized version of each signal, using a robust per-epoch approach based on the median (in place of the mean) and a robust estimate of the standard deviation (0.7413 times the interquartile range). For each epoch, we then computed an initial round of “level-1” features per individual: absolute spectral power (0.75 to 25 Hz in 0.25 Hz bins) and relative spectral power (as above, but normalized by total power in that frequency range) for both the original and time-domain normalized signals. For the time-domain normalized signals, we further computed a set of time-domain metrics: three Hjorth parameters and permutation entropy and fractal dimension [32]. We removed epochs with any time-domain metrics that had extreme outlier values (above 10 SD units from the mean for that individual) and then saved all level-1 epoch-wise metrics (computed independently per individual) in intermediate compact binary files.

We next concatenated all level-1 feature sets to construct a single multi-individual (“level-2”) training feature dataset, to a) derive principal components across feature sets as new features, b) derive further new features on-the-fly (e.g. temporally smoothed versions of original features) and then, c) to train the classifier. Specifically, we derived principal components by first mean-centering features within each individual, and then applying a singular value decomposition to extract K orthogonal components to replace a set of features. In particular, we extracted K=6 components from each absolute power spectra for the original and time-domain normalized signals, and K=4 components from the analogous relative power spectra. We speculated that including features that are normalized and unnormalized in both the time and frequency domains might improve the robustness of the model and make it more transferable to new datasets. Then for each set of principal components and the time-domain features (Hjorth, entropy and fractal dimension), we created new features that were temporally smoothed (within individual) versions of each, using a triangular moving average with a window of either 2, 10 or 25 minutes, to capture both abrupt and longer-term ultradian changes in the sleep EEG. We finally created normalized versions of the time-domain metrics and added a time-track feature (scaled 0 to 1 for each individual, increasing by 1/N_E_ units if that individual had N_E_ epochs). In total this yielded for the single-channel model 113 features representing each 30-second epoch in the final dataset. The majority of features therefore reflect principal spectral components of the EEG, with different forms of normalization and temporal smoothing applied.

#### Classification

Following [32], we employed the widely used Light Gradient-Boosting Machine (LightGBM, Ke et al 2017) framework to perform the classification task of predicting which of five sleep stages (N1, N2, N3, R and W) is most likely for each epoch. Airaclve aspects of this classifier include that it performs well in the presence of highly correlated features and allows for missing data.

Stages were assigned weights of 2.2, 1, 1.2, 1.4, and 1 to reflect the class imbalance in the original training data. When creating level-2 features and training, the singular vectors and values (W and V matrices) were saved and used to project the corresponding spectral metrics for each new individual/epoch into the previously defined principal component subspace.

Further, the mean and standard deviation of each metric were also saved and used at prediction: feature values that were more than 4 SD units from the mean were set to missing in each test individual; if more than 1/3 of all epochs had outlier values for a given feature/test individual, then all epoch for that feature were set to missing.

#### Pediatric models

Although it included individuals from the CHAT and CCSHS (adolescent) datasets, the original POPS model was trained on a sample containing a large proportion of older adults. Further, the original model was trained on single/central EEG channels, as those were typically the only channels available for many NSRR cohorts (e.g. SHHS). In the context of RASP, we reasoned that a stager trained on a larger number of pediatric samples might outperform the original POPS model. As the available pediatric datasets in NSRR typically had at least six EEG channels (i.e. frontal and occipital as well as central channels), we wanted to create a new staging model that also took advantage of these fuller montages. We followed a similar approach as for the adult model with the following exceptions. First, EEG data were summarized by 8 principal components based on spectra from (either lev or right) frontal, central and occipital channels jointly (as above, this was performed four times, with normalization in lme and/or frequency domains). Second, 4 principal components were derived from the EOG spectra under each condition. In total, this yielded 321 features per epoch that were passed to the LightGBM classifier under either model training or prediction. To disentangle the impact of a different training population from a different (expanded) feature set when comparing pediatric and adult stagers applied to the RASP cohorts, we further created a single-channel variant, using the same model as the adult stager (i.e. central channels, without EOG) but trained on the pediatric samples.

### EEG preprocessing

Sleep EEG data processing was performed using the open-source Luna toolset (<u>hip://zzz.bwh.harvard.edu/luna/</u>). All analyses focused on six EEG channels present in the RASP cohorts as well as the three NSRR cohorts (NCH-SDB, CHAT and PATS), namely F3, F4, C3, C4, O1, and O2, re-referenced to the contralateral mastoid (M1/M2) and resampled to 128 Hz. All signals were bandpass-filtered using a zero-phase Kaiser window finite impulse response filter, with transition frequencies at 0.3 and 45 Hz. Across either all NREM (N2 and N3) or all REM periods, we identified 30-second epochs that likely contained gross artifact, based on any of the six EEG channels having any of three Hjorth parameters (activity, mobility, complexity) [41] more than 4 standard deviation units above or below the mean for that individual/channel/stage; this procedure was performed twice, removing on average of 3.8% (12.72 minutes) of the NREM recording time per individual. Subsequent outlier removal was performed on a metric-by-metric basis for a given analysis, by removing individual’s whose scores were more than 3 SD units from the group mean.

### Sleep microarchitecture analysis

#### Spectral power, slope, and coherence

Following artifact removal, stage-specific spectral power was estimated using Welch’s method, using 4 second segments with 2 second overlaps (yielding a spectral resolution of 0.25 Hz) and applying a Tukey (cosine-tapered) window with α = 0.5, taking the median power across all segments in an epoch. Stage-specific power estimates were derived as the mean of the epoch-wise values. Band power values were expressed in log-scaled units (10log (*X*)), derived using the following definitions (in Hz): slow [0.5, 1), delta [1,4), theta [4,8), alpha [8,11), sigma [11,15) and beta [15,30). We estimated the 1/f spectral slope as previously described [42] based on a log-log linear regression of power in each 0.25 Hz bin from 30 to 45 Hz. To reduce the impact of artifact on this metric, we fit an initial model and removed data points (frequency-power pairs) that had residuals greater than 2 SD units from the mean when fixng the final model. To capture differences in functional connectivity between individuals, we computed the magnitude squared coherence between each pair of channels for each power band.

#### Spindle detection

We detected slow and fast spindles as previously described [43,44]. Briefly, we used a complex Morlet wavelet transform with center frequencies of 11 Hz and 15 Hz to target slow and fast spindles respectively and 7 cycles (corresponding to a detection window of approximately +/- 2 Hz). Spindle events were detected as intervals in which all wavelet coefficients were at least 4 times greater than the median for that individual/channel, for more than 0.5 but not more than 3 seconds. Spindles detected within 0.5 seconds of each other were merged if the resulting spindle was < 3 seconds.

We quantified spindle activity using the following metrics: density (spindles per minute), count, mean duration, mean amplitude (peak-to-peak, μV), mean integrated spindle activity (ISA) per spindle (ISA_S_), per minute (ISA_M_) and the total ISA (ISA_T_), mean number of oscillations per spindle, and secondary measures of spindle morphology, including mean chirp (intra-spindle change in frequency) and symmetry indices. The ISA is the area under the curve of the normalized wavelet power during spindle events, thereby reflecting both the duration and amplitude of the typical spindle (ISA_S_), and optionally, also the rate (ISA_M_) or the absolute number (ISA_T_) of spindle occurrences. Spindle chirp is the log of the ratio of mean peak-to-peak time intervals in the first versus the second half of the spindle. A first symmetry index, *S*, is the relative location of the spindle’s central peak (point of maximum peak-to-peak amplitude) scaled from 0 to 1; a second “folded” symmetry index is calculated as 2|*S*-0.5|.

#### Slow oscillations

We used an adaptive heuristic to detect slow oscillations (SO) as previously described [43]. Specifically, we band-pass filtered the EEG using transition frequencies of 0.3 and 4Hz, marked all positive to-negative zero-crossings, and designated SOs as those intervals between zero-crossings with a duration between 0.5 and 2 seconds (i.e. 0.5 to 2 Hz) and having both a peak-to-peak amplitude and an absolute minimum negative peak amplitude greater than twice the average value for that individual. We quantified SO activity using SO count and density (counts per minute) and SO morphology based on mean peak-to-peak amplitude (μV, log-scaled), duration or “wavelength” (seconds), and upward slope of the negative peak (μV/sec, log-scaled).

#### Spindle/SO coupling

We estimated three measures of coupling: gross overlap, coupling magnitude and coupling phase angle. Whereas ‘overlap’ measured whether spindles and SO tended to occur concurrently, ‘phase coupling’ measured whether spindles tended to peak at particular times during the SO, thereby indexing a more precise temporal relationship. All measures were defined relative to spindle peaks: the point of maximal peak-to-peak amplitude. Overlap was the number of spindle peaks that fell within a detected SO. We determined the null distribution for this metric empirically, by randomly shuffling spindle peaks and recalculating overlap 10,000 times. For each individual, we normalized the overlap metric as a Z score, given the mean and standard deviation from the null distribution. Coupling magnitude and phase angle metrics were based on the instantaneous phase from a Hilbert transform of EEG aver bandpass-filtering in the 0.3 to 4 Hz range. We calculated intra-trial phase consistency (ITPC) as a measure of the strength of spindle/SO coupling. If the Hilbert-derived phase angle at each spindle peak is assumed to be a unit vector on a circle, with an angle matching the phase angle, then the ITPC is the magnitude of the average of these complex vectors. An individual’s coupling phase angle is given by the angle of the complex mean of these vectors. As ITPC statistics and asymptotic p-values can show bias or noise when based on a small number of spindle/SO events, or on non-sinusoidal waveforms, we generated empirical null distributions to normalize them, based on shuffling spindle peaks 10,000 times. To control for gross spindle/SO overlap, we shuffled each spindle peak by a random offset, between o seconds and the duration of the spanning SO, wrapping as necessary. This ‘within-SO’ shuffling scheme preserved the total number of spindles, SO and their gross overlap in each null replicate, but randomized only the precise relationships between spindle peaks and SO phase.

### Feature selecDon for age predicDon model

To identify measures to include in our model, we used pediatric cases without NDDs in NCH-SDB, CHAT, and PATS. We performed linear regressions on 1338 individual Z-scored microarchitectural sleep features on age, correcting for sex, race, and study site. These features represented 66 unique metrics, repeated for paired and averaged frontal, central, and occipital channels, seven band frequencies, and three sleep stages (N2, N3, and REM). Sleep features that went into the final predictive model explained more than 2.5% of the variance in age. We excluded features that showed inter-site variability, identifying these by comparing the significance values of the age covariate with site covariates. Metrics with a -log_10_(p-value) of the age covariate less than three times that of the maximum -log10(p-value) of the site covariates were excluded. In all, our initial feature set included 448 metrics. We removed highly correlated features (r > 0.7) and performed Lasso regression with the glmnet package in R for variable selection. As a final step, we utilized forward and backwards stepwise model selection by AIC. For models with restricted feature subsets, we repeated the above steps but started with features from specific subsets (for example, features from only N2 sleep stage).

### StaDsDcal analysis

All stalslcal analyses were performed using R version 4.3.1.

#### Pediatric versus automated stager performance

Automated stager performance was assessed by comparing manual versus automated staging kappa and accuracy at transitions. To assess transition accuracies, we focused on epochs flanked by two of the same stages and on transitions between two epochs of different stages. Multiple paired T-tests with Bonferroni correction were used to compare across the three automated stagers.

#### Correlating manual versus automated staging sleep architecture

To compare sleep architectural features derived from manual versus automated staging in the RASP cohorts, we focused on macroarchitectural and overall NREM metrics (N2 and N3 combined). Pearson’s correlation coefficient was calculated for 1828 total features, comprising 61 metrics spanning six domains (macroarchitecture, spectral band power, spectral slope, slow oscillations, spindles, and interchannel connectivity), repeated for 10 EEG channels, seven band frequencies, and slow (11 Hz) and fast (15 Hz) spindle frequencies. To assess whether developmental changes in sleep architecture were conserved between manual and automated staging, we regressed individual Z-scored sleep features on age, correcting for sex, diagnostic status, and study site. We calculated the Pearson’s correlation coefficient between the beta of the age covariate derived from manual versus automated staging. To determine whether automated staging conserves diagnostic changes in sleep architecture, we regressed Z-scored individual sleep metrics on diagnosis, correcting for age, sex, and study site. We compared the resultant differences between non-NDD and NDD groups (i.e., the beta of the diagnosis covariate) across manual versus automated staging with Pearson’s correlation coefficient.

#### Comparing sleep architecture across diagnostic status and cohorts

We compared individual sleep features between non-NDD and DS or ASD cases in both RASP and NCH-SDB cohorts. We examined 1351 total sleep features across 79 metrics in the six domains (see above). Each metric was repeated for paired and averaged frontal, central, and occipital channels, seven band frequencies, and three sleep stages (N2, N3, and REM). We performed logistic regressions of individual Z-scored sleep features on diagnostic status and reported the beta of the diagnosis covariate (i.e., the standardized difference in sleep feature between diagnostic status).

#### Assessing age prediction model performance

To compare model performance across subsets of sleep features, we performed repeated k-fold cross validation (3 repeats and 10 fold) with the NCH-SDB, PATS, and CHAT non-NDD cases. We compared MAE and R^2^ across all 30 folds between our original full feature model and each specific subset model, as well as between our random subset feature model and each specific subset model using multiple T-tests with Bonferroni correction.

#### Age prediction using sleep architecture features

We predicted age in the RASP cohorts as well as in an NCH-SDB hold-out set (30% of the non-NDD NCH-SDB cases). The remaining 70% of the NCH-SDB non-NDD cases, as well as all of PATS and CHAT cohorts were used as a training set with the final model features (see above: “Feature selection for age prediction model”). We also predicted age in DS and ASD cases from both the RASP as well as the NCH-SDB cohorts. We predicted age 100x, resampling the NCH-SDB non-NDD test set each time while holding the RASP and NDD cohorts constant. For each resample, we compared MAE and ME (i.e., predicted age gap) between non-NDD and overall NDD, DS, or ASD cohorts in NCH-SDB or RASP, correcting for age and sex, and additionally study site in RASP.

#### Comparing variances between diagnostic groups

Age- and sex-matched non-NDD controls were selected to compare against RASP NDD ASD or DS cases. To decrease the skewness of absolute band power data, we compared log10-transformed absolute power values. To account for differences arising from study sites rather than diagnostic status, we took the following steps for each individual sleep metric. First, outliers (defined by >3 standard deviations from the mean of the entire group) were removed. Next, we regressed each individual metric on site, correcting for age, sex, and diagnostic status and obtained the residuals of the fitted model to adjust for site effects. Finally, we removed outliers from residualized sleep metrics and compared variances across diagnostic groups using the Fligner-Killeen test. P-values were FDR-adjusted to account for multiple comparisons.

## Acknowledgements

This work was partially supported by the Pediatric Epilepsy Research Foundation (to M.M.S. and A.B.).

## References

1. Roffwarg HP, Muzio JN, Dement WC. Ontogenetic development of the human sleep-dream cycle. Science. 1966;152(3722):604–619. doi:10.1126/science.152.3722.604

2. Weissbluth M. Naps in children: 6 months-7 years. Sleep. 1995;18(2):82–87. doi:10.1093/sleep/18.2.82

3. Iglowstein I, Jenni OG, Molinari L, Largo RH. Sleep duration from infancy to adolescence: reference values and generational trends. Pediatrics. 2003;111(2):302–307. doi:10.1542/peds.111.2.302

4. Baker FC, Willoughby AR, Massimiliano de Z, et al. Age-Related Differences in Sleep Architecture and Electroencephalogram in Adolescents in the National Consortium on Alcohol and Neurodevelopment in Adolescence Sample. Sleep. 2016;39(7):1429–1439. doi:10.5665/sleep.5978

5. Feinberg I, Davis NM, de Bie E, Grimm KJ, Campbell IG. The maturational trajectories of NREM and REM sleep durations differ across adolescence on both school-night and extended sleep. Am J Physiol-Regul Integr Comp Physiol. 2012;302(5):R533–R540. doi:10.1152/ajpregu.00532.2011

6. Bridi MCD, Aton SJ, Seibt J, Renouard L, Coleman T, Frank MG. Rapid eye movement sleep promotes cortical plasticity in the developing brain. Sci Adv. 2015;1(6):e1500105. doi:10.1126/sciadv.1500105

7. Cao J, Herman AB, West GB, Poe G, Savage VM. Unraveling why we sleep: Quantitative analysis reveals abrupt transition from neural reorganization to repair in early development. Sci Adv. 2020;6(38):eaba0398. doi:10.1126/sciadv.aba0398

8. Frank MG, Issa NP, Stryker MP. Sleep enhances plasticity in the developing visual cortex. Neuron. 2001;30(1):275–287. https://www.ncbi.nlm.nih.gov/pubmed/11343661

9. Kayser MS, Yue Z, Sehgal A. A critical period of sleep for development of courtship circuitry and behavior in Drosophila. Science. 2014;344(6181):269–274. doi:10.1126/science.1250553 [doi]

10. Knoop MS, Groot ER de, Dudink J. Current ideas about the roles of rapid eye movement and non–rapid eye movement sleep in brain development. Acta Paediatr. 2021;110(1):36–44. doi:10.1111/apa.15485

11. Sokoloff G, Dooley JC, Glanz RM, et al. Twitches emerge postnatally during quiet sleep in human infants and are synchronized with sleep spindles. Curr Biol. Published online June 17, 2021. doi:10.1016/j.cub.2021.05.038

12. Buxton OM, Chang AM, Spilsbury JC, Bos T, Emsellem H, Knutson KL. Sleep in the modern family: protective family routines for child and adolescent sleep. Sleep Health. 2015;1(1):15–27. doi:10.1016/j.sleh.2014.12.002

13. Friel CP, Duran AT, Shechter A, Diaz KM. U.S. Children Meeting Physical Activity, Screen Time, and Sleep Guidelines. Am J Prev Med. 2020;59(4):513–521. doi:10.1016/j.amepre.2020.05.007

14. James S, Chang AM, Buxton OM, Hale L. Disparities in adolescent sleep health by sex and ethnoracial group. SSM - Popul Health. 2020;11:100581. doi:10.1016/j.ssmph.2020.100581

15. Wheaton AG, Jones SE, Cooper AC, Crov JB. Short Sleep Duration Among Middle School and High School Students — United States, 2015. Morb Mortal Wkly Rep. 2018;67(3):85–90. doi:10.15585/mmwr.mm6703a1

16. Beebe DW. Cognitive, Behavioral, and Functional Consequences of Inadequate Sleep in Children and Adolescents. Pediatr Clin North Am. 2011;58(3):649–665. doi:10.1016/j.pcl.2011.03.002

17. Booth SA, Carskadon MA, Young R, Short MA. Sleep duration and mood in adolescents: an experimental study. Sleep. 2021;44(5):zsaa253. doi:10.1093/sleep/zsaa253

18. Talbot LS, McGlinchey EL, Kaplan KA, Dahl RE, Harvey AG. Sleep deprivation in adolescents and adults: changes in affect. Emot Wash DC. 2010;10(6):831–841. doi:10.1037/a0020138

19. Huhdanpää H, Morales-Muñoz I, Aronen ET, et al. Sleep Difficulties in Infancy Are Associated with Symptoms of Inaienlon and Hyperactivity at the Age of 5 Years: A Longitudinal Study. J Dev Behav Pediatr. 2019;40(6):432. doi:10.1097/DBP.0000000000000684

20. Morales-Muñoz I, Nolvi S, Mäkelä T, et al. Sleep during infancy, inhibitory control and working memory in toddlers: findings from the FinnBrain cohort study. Sleep Sci Pract. 2021;5(1):13. doi:10.1186/s41606-021-00064-4

21. Sivertsen B, Harvey AG, Reichborn-Kjennerud T, Torgersen L, Ystrom E, Hysing M. Later emotional and behavioral problems associated with sleep problems in toddlers: a longitudinal study. JAMA Pediatr. 2015;169(6):575–582. doi:10.1001/jamapediatrics.2015.0187

22. Angriman M, Caravale B, Novelli L, Ferri R, Bruni O. Sleep in Children with Neurodevelopmental Disabilities. Neuropediatrics. 2015;46(3):199–210. doi:10.1055/s-0035-1550151

23. Kamara D, Beauchaine TP. A Review of Sleep Disturbances among Infants and Children with Neurodevelopmental Disorders. Rev J AuDsm Dev Disord. 2020;7(3):278–294. doi:10.1007/s40489-019-00193-8

24. Robinson-Shelton A, Malow BA. Sleep Disturbances in Neurodevelopmental Disorders. Curr Psychiatry Rep. 2016;18(1):6. doi:10.1007/s11920-015-0638-1

25. Veatch OJ, Sutcliffe JS, Warren ZE, Keenan BT, Potter MH, Malow BA. Shorter sleep duration is associated with social impairment and comorbidities in ASD. AuDsm Res Off J Int Soc AuDsm Res. 2017;10(7):1221–1238. doi:10.1002/aur.1765

26. Reynolds AM, Spaeth AM, Hale L, et al. Pediatric sleep: current knowledge, gaps, and opportunities for the future. Sleep. 2023;46(7):zsad060. doi:10.1093/sleep/zsad060

27. Fernandez LMJ, Lüthi A. Sleep Spindles: Mechanisms and Functions. Physiol Rev. 2020;100(2):805–868. doi:10.1152/physrev.00042.2018

28. Kozhemiako N, Buckley AW, Chervin RD, Redline S, Purcell SM. Mapping neurodevelopment with sleep macro- and micro-architecture across multiple pediatric populations. NeuroImage Clin. 2024;41:103552. doi:10.1016/j.nicl.2023.103552

29. Moser D, Anderer P, Gruber G, et al. Sleep classification according to AASM and Rechtschaffen & Kales: effects on sleep scoring parameters. Sleep. 2009;32(2):139–149. doi:10.1093/sleep/32.2.139

30. Grigg-Damberger MM. The Visual Scoring of Sleep in Infants 0 to 2 Months of Age. J Clin Sleep Med JCSM Off Publ Am Acad Sleep Med. 2016;12(3):429–445. doi:10.5664/jcsm.5600

31. Perslev M, Darkner S, Kempfner L, Nikolic M, Jennum PJ, Igel C. U-Sleep: resilient high-frequency sleep staging. Npj Digit Med. 2021;4(1):1–12. doi:10.1038/s41746-021-00440-5

32. Vallat R, Walker MP. An open-source, high-performance tool for automated sleep staging. Peyrache A, Büchel C, Bagur S, eds. eLife. 2021;10:e70092. doi:10.7554/eLife.70092

33. Baumert M, Hartmann S, Phan H. Automatic sleep staging for the young and the old – Evaluating age bias in deep learning. Sleep Med. 2023;107:18–25. doi:10.1016/j.sleep.2023.04.002

34. Sun H, Paixao L, Oliva JT, et al. Brain age from the electroencephalogram of sleep. Neurobiol Aging. 2019;74:112–120. doi:10.1016/j.neurobiolaging.2018.10.016

35. Kozhemiako N, Jiang C, Sun Y, et al. A spectrum of altered non-rapid eye movement sleep in schizophrenia. Published online December 29, 2023:2023.12.28.573548. doi:10.1101/2023.12.28.573548

36. Zhang Y, Kim M, Prerau M, et al. The National Sleep Research Resource: making data findable, accessible, interoperable, reusable and promoting sleep science. Sleep. 2024;47(7):zsae088. doi:10.1093/sleep/zsae088

37. Lee H, Li B, DeForte S, et al. A large collection of real-world pediatric sleep studies. Sci Data. 2022;9(1):421. doi:10.1038/s41597-022-01545-6

38. Marcus CL, Moore RH, Rosen CL, et al. A randomized trial of adenotonsillectomy for childhood sleep apnea. N Engl J Med. 2013;368(25):2366–2376. doi:10.1056/NEJMoa1215881

39. Redline S, Cook K, Chervin RD, et al. Adenotonsillectomy for Snoring and Mild Sleep Apnea in Children: A Randomized Clinical Trial. JAMA. 2023;330(21):2084–2095. doi:10.1001/jama.2023.22114

40. Wang R, Bakker JP, Chervin RD, et al. Pediatric Adenotonsillectomy Trial for Snoring (PATS): protocol for a randomised controlled trial to evaluate the effect of adenotonsillectomy in treating mild obstructive sleep-disordered breathing. BMJ Open. 2020;10(3):e033889. doi:10.1136/bmjopen-2019-033889

41. Hjorth B. EEG analysis based on lme domain properties. Electroencephalogr Clin Neurophysiol. 1970;29(3):306–310. doi:10.1016/0013-4694(70)90143-4

42. Kozhemiako N, Mylonas D, Pan JQ, Prerau MJ, Redline S, Purcell SM. Sources of Variation in the Spectral Slope of the Sleep EEG. eNeuro. 2022;9(5). doi:10.1523/ENEURO.0094-22.2022

43. Djonlagic I, Mariani S, Fitzpatrick AL, et al. Macro and micro sleep architecture and cognitive performance in older adults. Nat Hum Behav. 2021;5(1):123–145. doi:10.1038/s41562-020-00964-y

44. Purcell SM, Manoach DS, Demanuele C, et al. Characterizing sleep spindles in 11,630 individuals from the National Sleep Research Resource. Nat Commun. 2017;8:15930. doi:10.1038/ncomms15930

45. Danker-Hopfe H, Kunz D, Gruber G, et al. Interrater reliability between scorers from eight European sleep laboratories in subjects with different sleep disorders. J Sleep Res. 2004;13(1):63–69. doi:10.1046/j.1365-2869.2003.00375.x

46. Magalang UJ, Chen NH, Cistulli PA, et al. Agreement in the scoring of respiratory events and sleep among international sleep centers. Sleep. 2013;36(4):591–596. doi:10.5665/sleep.2552

47. Rosenberg RS, Van Hout S. The American Academy of Sleep Medicine inter-scorer reliability program: sleep stage scoring. J Clin Sleep Med JCSM Off Publ Am Acad Sleep Med. 2013;9(1):81–87. doi:10.5664/jcsm.2350

48. Younes M, Raneri J, Hanly P. Staging Sleep in Polysomnograms: Analysis of Inter-Scorer Variability. J Clin Sleep Med JCSM Off Publ Am Acad Sleep Med. 2016;12(6):885–894. doi:10.5664/jcsm.5894

49. Zhang X, Dong X, Kantelhardt JW, et al. Process and outcome for international reliability in sleep scoring. Sleep Breath Schlaf Atm. 2015;19(1):191–195. doi:10.1007/s11325-014-0990-0

50. Saeys Y, Inza I, Larrañaga P. A review of feature selection techniques in bioinformatics. BioinformaDcs. 2007;23(19):2507–2517. doi:10.1093/bioinformatics/btm344

51. Simpson R, Oyekan AA, Ehsan Z, Ingram DG. Obstructive sleep apnea in patients with Down syndrome: current perspectives. Nat Sci Sleep. 2018;10:287–293. doi:10.2147/NSS.S154723

52. Durdik P, Sujanska A, Suroviakova S, Evangelisti M, Banovcin P, Villa MP. Sleep Architecture in Children With Common Phenotype of Obstructive Sleep Apnea. J Clin Sleep Med JCSM Off Publ Am Acad Sleep Med. 2018;14(1):9–14. doi:10.5664/jcsm.6868

53. D’Rozario AL, Cross NE, Vakulin A, et al. Quantitative electroencephalogram measures in adult obstructive sleep apnea – Potential biomarkers of neurobehavioural functioning. Sleep Med Rev. 2017;36:29–42. doi:10.1016/j.smrv.2016.10.003

54. Parker JL, Melaku YA, D’Rozario AL, et al. The association between obstructive sleep apnea and sleep spindles in middle-aged and older men: a community-based cohort study. Sleep. 2022;45(3):zsab282. doi:10.1093/sleep/zsab282

55. Cohen S, Fulcher BD, Rajaratnam SMW, et al. Behaviorally-determined sleep phenotypes are robustly associated with adaptive functioning in individuals with low functioning autism. Sci Rep. 2017;7(1):1–8. doi:10.1038/s41598-017-14611-6

56. Mazzone L, Postorino V, Siracusano M, Riccioni A, Curatolo P. The Relationship between Sleep Problems, Neurobiological Alterations, Core Symptoms of Autism Spectrum Disorder, and Psychiatric Comorbidities. J Clin Med. 2018;7(5):102. doi:10.3390/jcm7050102

57. Touchette E, Petit D, Seguin JR, Boivin M, Tremblay RE, Montplaisir JY. Associations between sleep duration patterns and behavioral/cognitive functioning at school entry. Sleep. 2007;30(9):1213–1219. https://www.ncbi.nlm.nih.gov/pubmed/17910393

58. MacDuffie KE, Munson J, Greenson J, et al. Sleep Problems and Trajectories of Restricted and Repetitive Behaviors in Children with Neurodevelopmental Disabilities. J AuDsm Dev Disord. 2020;50(11):3844–3856. doi:10.1007/s10803-020-04438-y

59. MacDuffie KE, Shen MD, Dager SR, et al. Sleep Onset Problems and Subcortical Development in Infants Later Diagnosed With Autism Spectrum Disorder. Am J Psychiatry. 2020;177(6):518–525. doi:10.1176/appi.ajp.2019.19060666

60. Coll-Tané M, Gong NN, Belfer SJ, et al. The CHD8/CHD7/Kismet family links blood-brain barrier glia and serotonin to ASD-associated sleep defects. Sci Adv. 2021;7(23):eabe2626. doi:10.1126/sciadv.abe2626

61. Gong NN, Dilley LC, Williams CE, et al. The chromatin remodeler ISWI acts during Drosophila development to regulate adult sleep. Sci Adv. 2021;7(8):eabe2597. doi:10.1126/sciadv.abe2597

62. Medina E, Schoch H, Ford K, Wintler T, Singletary KG, Peixoto L. Shank3 influences mammalian sleep development. J Neurosci Res. 2022;100(12):2174–2186. doi:10.1002/jnr.25119

63. Williams Buckley A, Hirtz D, Oskoui M, et al. Practice guideline: Treatment for insomnia and disrupted sleep behavior in children and adolescents with autism spectrum disorder. Neurology. 2020;94(9):392–404. doi:10.1212/WNL.0000000000009033

64. Cole JH, Annus T, Wilson LR, et al. Brain-predicted age in Down syndrome is associated with beta amyloid deposition and cognitive decline. Neurobiol Aging. 2017;56:41–49. doi:10.1016/j.neurobiolaging.2017.04.006

65. Horvath S, Garagnani P, Bacalini MG, et al. Accelerated epigenetic aging in Down syndrome. Aging Cell. 2015;14(3):491–495. doi:10.1111/acel.12325

66. Xu K, Li S, Muskens IS, et al. Accelerated epigenetic aging in newborns with Down syndrome. Aging Cell. 2022;21(7):e13652. doi:10.1111/acel.13652

67. Gameren-Oosterom HBM van, Fekkes M, Buitendijk SE, Mohangoo AD, Bruil J, Wouwe JPV. Development, Problem Behavior, and Quality of Life in a Population Based Sample of Eight-Year-Old Children with Down Syndrome. PLOS ONE. 2011;6(7):e21879. doi:10.1371/journal.pone.0021879

68. Banville H, Jaoude MA, Wood SUN, et al. Do try this at home: Age prediction from sleep and meditation with large-scale low-cost mobile EEG. Imaging Neurosci. 2024;2:1–15. doi:10.1162/imag_a_00189

69. Tetereva A, Pat N. The (Limited?) Utility of Brain Age as a Biomarker for Capturing Fluid Cognition in Older Individuals. eLife. 2023;12. doi:10.7554/eLife.87297.2

