## Supplementary Figures for "Leveraging clinical sleep data across multiple pediatric cohorts for insights into neurodevelopment: the Retrospective Analysis of Sleep in Pediatric (RASP) cohorts study"

### Supplementary Materials

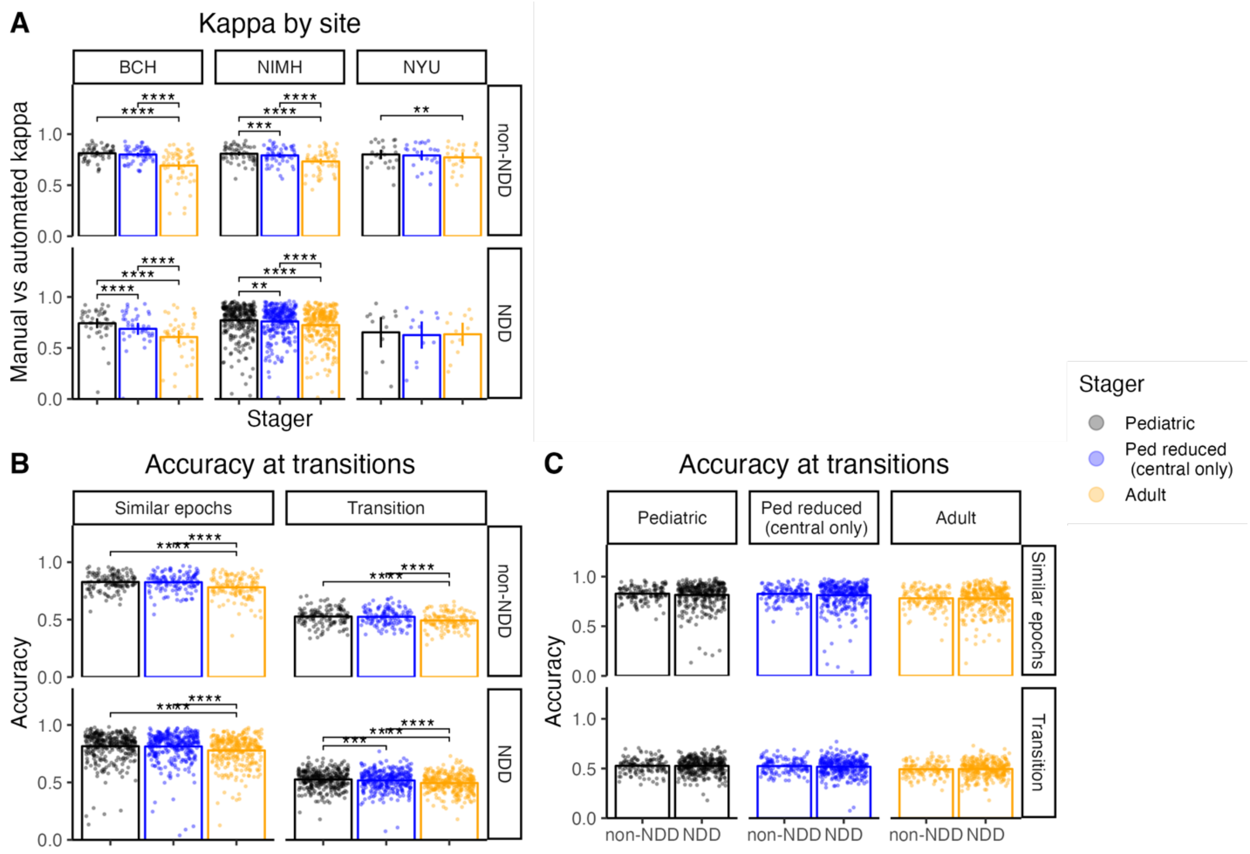


**Figure S1: Automated stager performance in specific RASP cohorts and at transitions.** A) Kappa for 3-class staging between observed vs predicted stage by epoch per individual in pediatric stagers (black and blue) and adult stager (yellow), grouped by study site and NDD diagnosis. B) Accuracy at transitions predicted by automated stagers between two epochs of the same (left) vs different (right) stages, grouped by NDD diagnosis. C) Comparison of accuracy at transitions predicted by automated stagers between two epochs of the same (top) vs different (bottom) stages between typical and NDD cases. Statistical comparisons in A and B were made using multiple paired T-tests with Bonferroni correction; comparisons in C were made using logistic regression correcting for age, sex, and study site (* p<0.05, ** p < 0.01, *** p < 0.001).

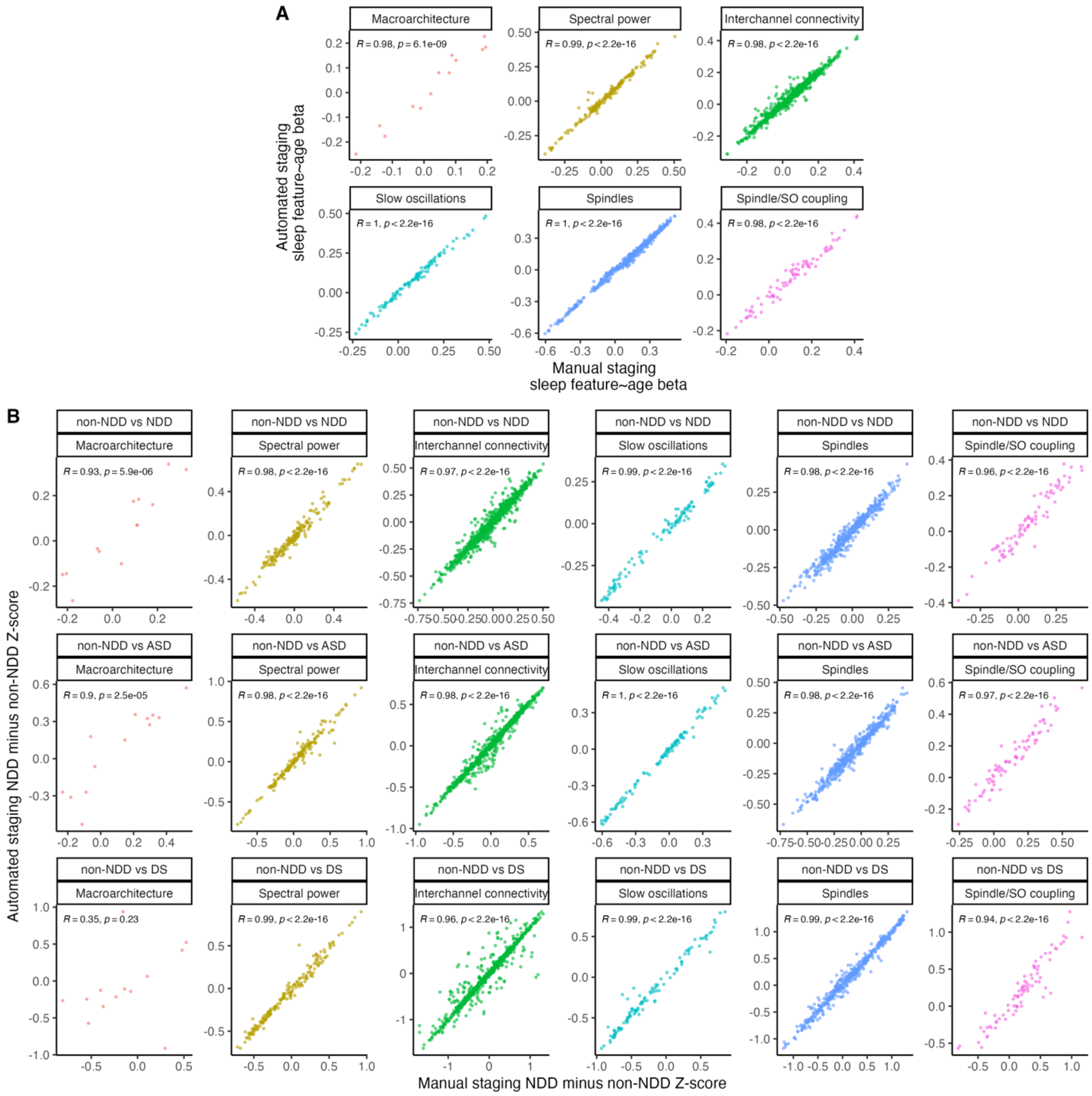


**Figure S2: Developmental and diagnostic changes in sleep metrics, by sleep metric class.** A) Developmental changes in sleep architecture metrics by class, comparing features derived from manual versus automated staging. B) Diagnostic changes in sleep architecture metrics by class and diagnosis, comparing features derived from manual versus automated staging. For both A and B, each point represents a sleep metric. X and Y axes denote the beta of the age and diagnosis coefficient, respectively, in the regression equation. R denotes Pearson's correlation coefficient and associated p-value is shown.

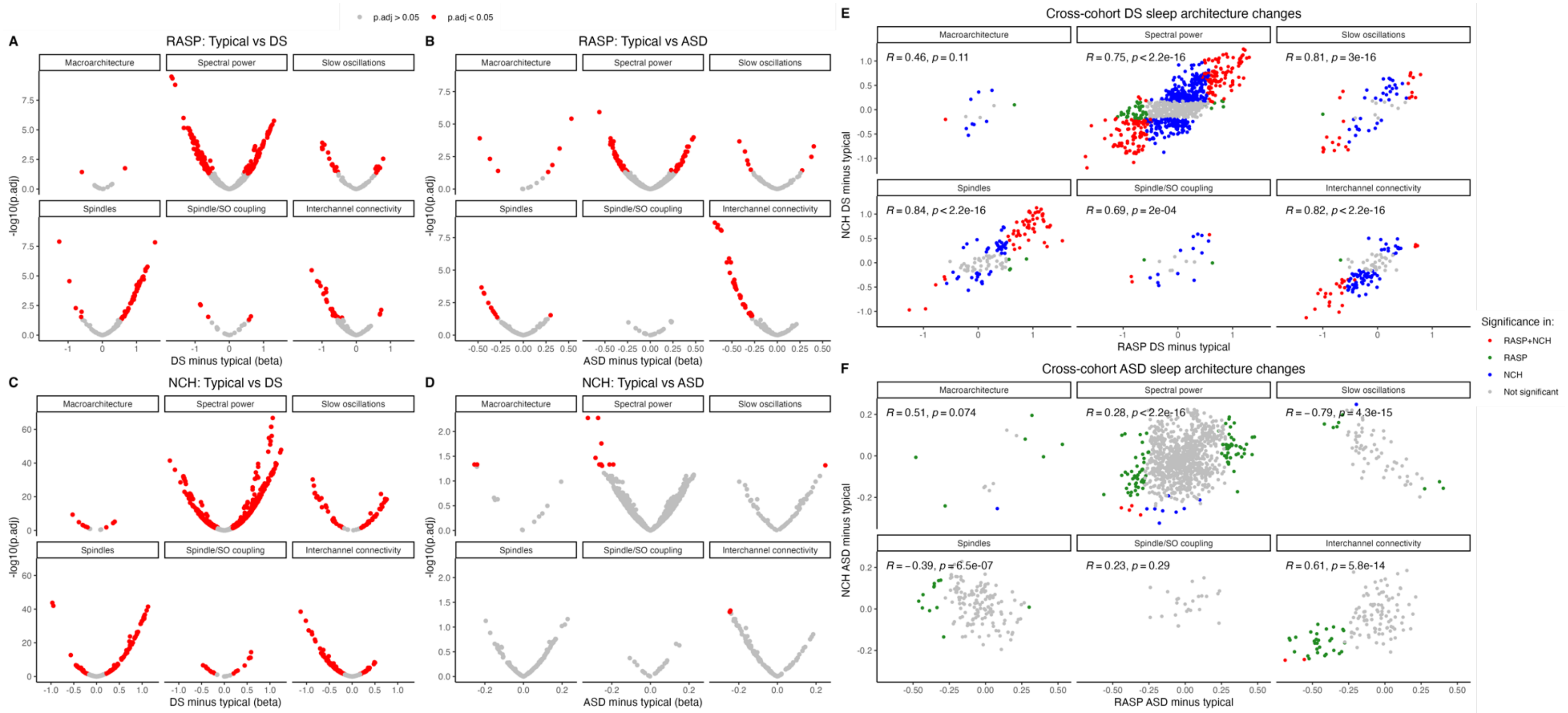


**Figure S4: Consistent sleep architecture changes in DS but not ASD, by domain.** Difference between denoted diagnosis and typical development versus –log10(p-value) of difference in A-B) RASP and C-D) NCH cohorts. E, F) Cross-cohort comparison of sleep architecture metrics in DS and ASD (respectively) vs typical development by sleep domain.

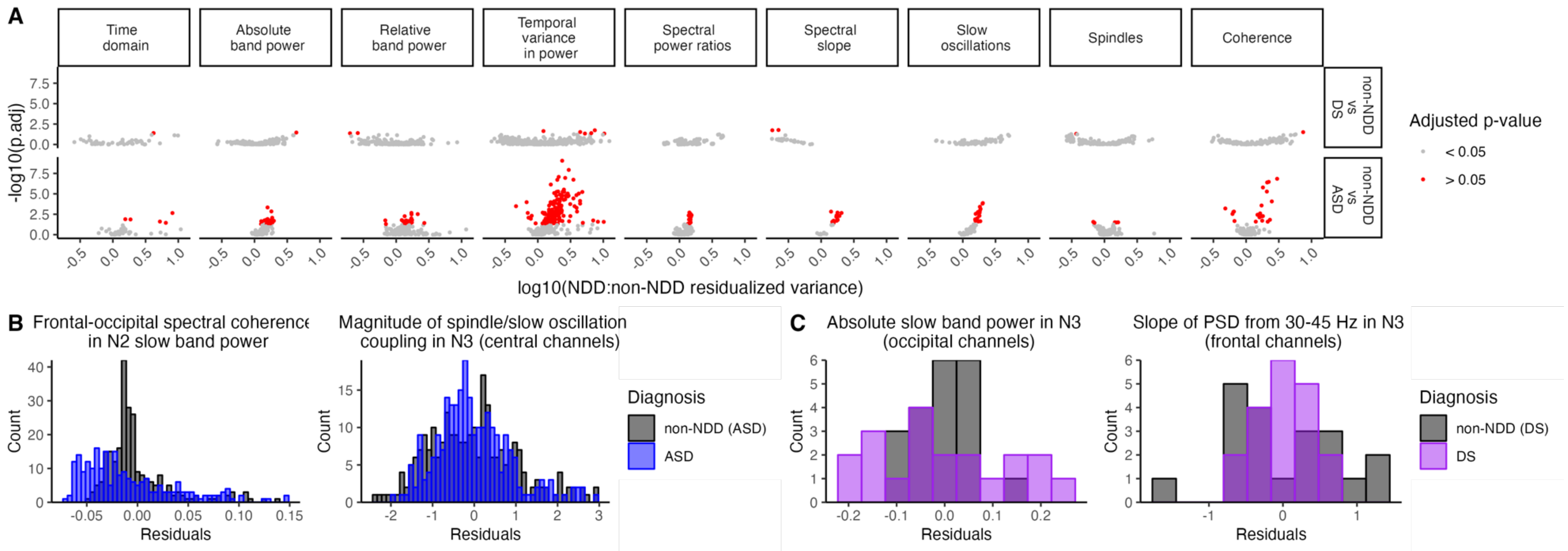


**Figure S5: Variance across individual sleep metrics from RASP ASD and DS cases.** A) log10 of the ratio of individual sleep metric variances between NDD vs non-NDD cases versus –log10 of the FDR-adjusted p-value comparing site-corrected variances between DS or ASD and age- and sex-matched non-NDD cases. Each individual point is a sleep metric. Metrics with significantly different variances across the denoted groups are in red. Examples of metrics with increased (left) and decreased (right) variances in B) ASD (blue) and C) DS (purple) compared to non-NDD cases (black). For all graphs in this figure, an equivalent number of age- and sex-matched non-NDD controls from the RASP cohort were used to compare variances of non-NDD vs NDD groups. For A, Bartlett's test was used to compare variances of individual metrics in the matched non-NDD and NDD cohorts.
